# Fast and accurate assembly of Nanopore reads via progressive error correction and adaptive read selection

**DOI:** 10.1101/2020.02.01.930107

**Authors:** Ying Chen, Fan Nie, Shang-Qian Xie, Ying-Feng Zheng, Thomas Bray, Qi Dai, Yao-Xin Wang, Jian-feng Xing, Zhi-Jian Huang, De-Peng Wang, Li-Juan He, Feng Luo, Jian-Xin Wang, Yi-Zhi Liu, Chuan-Le Xiao

**Author notes:** To whom correspondence should be addressed: Feng Luo.; Jian-Xing Wang.; Yi-Zhi Liu.; Chuan-Le Xiao. These authors contributed equally to the manuscript as first authors (Ying Chen, Fan Nie, Shang-Qian Xie, Ying-Feng Zheng).

## Abstract

Although long Nanopore reads are advantageous in *de novo* genome assembly, applying Nanopore reads in genomic studies is still hindered by their complex errors. Here, we developed NECAT, an error correction and *de novo* assembly tool designed to overcome complex errors in Nanopore reads. We proposed an adaptive read selection and two-step progressive method to quickly correct Nanopore reads to high accuracy. We introduced a two-stage assembler to utilize the full length of Nanopore reads. NECAT achieves superior performance in both error correction and *de novo* assembly of Nanopore reads. NECAT requires only 7,225 CPU hours to assemble a 35X coverage human genome and achieves a 2.28-fold improvement in NG50. Furthermore, our assembly of the human WERI cell line showed an NG50 of 29 Mbp. The high-quality assembly of Nanopore reads can significantly reduce false positives in structure variation detection.

---

Reconstructing the genome sequence of a species or individual in a population is one of the most important tasks in genomics^1-3^. Single-molecule sequencing (SMS) technologies, developed by Pacific Bioscience and Oxford Nanopore, yield long reads that can significantly increase the number of solvable repetitive genome regions and improve the contiguity of assembly^4-7^. However, SMS reads usually have high error rates^8^. The two strategies currently used for *de-novo* genome assembly from SMS reads are “correction then assembly” and “assembly then correction.” Assemblers, such as Falcon^9^, Canu^10^, and MECAT^11^, first correct SMS reads and then assemble the genome using corrected reads. Conversely, assemblers, such as miniasm^12^, Flye^13^ and wtdbg2^14^, assemble the genome using error-prone reads and then correct the assembled genome. Due to high computational cost of error correction, the “correction then assembly” approach is usually slower than “assembly then correction”. However, directly assembling the genome using error-prone SMS reads can increase assembly errors in the genome sequence, which affects the quality of reference genome and results in bias in downstream analysis, especially in complicated genome regions^10, 15^. On the other hand, the “correction then assembly” approach can provide highly continuous and accurate genome assemblies^9-11^.

The recently released R9 flow cell from Oxford Nanopore technology can generate reads that are up to 1M in length and with read N50 >100 kb, which may significantly improve the contiguity of assembly compared with those of assemblies using PacBio SMRT reads^5-7, 16^. However, errors in Nanopore reads are more complex than those in PacBio reads^17, 18^ (see Results). Error correction tools in current assemblers were originally designed for PacBio SMRT reads and cannot correct Nanopore reads efficiently and effectively. For example, correcting 30X coverage human Nanopore reads using error correction tool in Canu requires 29K CPU hours^16^. Moreover, the average identity of reads corrected by Canu is only 92%, which is far less accurate than that of corrected PacBio SMRT reads. These high error rates in corrected Nanopore reads can introduce mis-assemblies. Furthermore, high-error-rate subsequences in Nanopore reads are usually trimmed during error correction, which reduces both the length of original reads and contiguity of final assembly.

In this study, we developed NECAT, a novel error correction and *de novo* assembly tool designed to overcome the problem of complex errors in Nanopore reads. Unlike existing error correction tools that iteratively correct Nanopore reads, we developed a two-step progressive method for Nanopore-read correction. In the first step, NECAT corrects low error rate subsequences (LERS), while in the second step, it corrects high error rate subsequences (HERS), of the read. This progressive approach allows NECAT to quickly correct Nanopore reads, resulting in high accuracy of corrected reads. To fully take advantage of Nanopore-read length, we presented a two-stage assembler in NECAT. This assembler constructs contigs using corrected Nanopore reads, and then bridges the contigs using original raw reads. We also used an adaptive selection mechanism to choose high-quality supporting reads for each template read during error correction, and to select high-quality overlaps for each read during the read-overlap step. Our results indicate that NECAT achieves superior performance in error correction and *de novo* assembly of Nanopore reads.

## Results

### Analysis of sequencing errors in Nanopore reads

We analyzed sequencing errors in Nanopore reads of *E. coli, S. cerevisiae, A. thaliana, D. melanogaster, C. reinhardtii, O. sativa, S. pennellii* and *H. sapiens* (NA12878) (**Supplementary Note 1-5 and Supplementary Table 1-2**). As shown in **Supplementary Table 3**, average error rates of Nanopore reads for these eight species ranged from 12% (for *S. cerevisiae*) to 20.1% (for *A. thaliana*). Although average error rates of Nanopore reads are similar to those of PacBio SMRT reads, error rates in Nanopore reads are more broadly distributed than those of PacBio SMRT reads. The error rates of raw reads in the eight datasets used in our study were broadly distributed between 7-50% and centralized between 10-30% (**Figure 1A**).

**Table 1.** Performance comparison of Nanopore read error correction

| Datasets | Pipeline | Size(g)/Time(h)<br>/Speed(g/h) | Error<br>rate(%) | <=5%(%) | N50 | N75 | Read number<br>with HERS |
| --- | --- | --- | --- | --- | --- | --- | --- |
| E.coli | raw reads | 1.38/--/-- | 17.8 | 0.01 | 41,074 | 35,484 | 121 |
|  | Canu | 0.22/1.63/0.14 | 7.06 | 20.45 | 37,747 | 32,127 | 1 |
|  | NECAT | 1.41/0.76/1.86 | 2.23 (4.27) | 99.34(80.51) | 43,140 | 37,502 | 1 |
| S. cerevisiae | raw reads | 5.48/--/-- | 12 | 1.61 | 34,668 | 28,152 | 7,589 |
|  | Canu | 2.18/30.83/0.071 | 3.13 | 87.3 | 10,554 | 4,567 | 4,820 |
|  | NECAT | 4.57/3.90/1.17 | 1.53 (3.08) | 95.04(88.09) | 31,364 | 24,480 | 268 |
| D. melanogaster | raw reads | 8.30/--/-- | 16.2 | 2.3 | 17,730 | 13,621 | 12,438 |
|  | Canu | 4.79/18.10/0.26 | 8.15 | 57.57 | 15,220 | 10,658 | 6,523 |
|  | NECAT | 7.52/4.20/1.79 | 4.89 (7.03) | 72.03(64.18) | 17,369 | 13,104 | 3,481 |
| A. thaliana | raw reads | 3.08/--/-- | 20.1 | 1.57 | 23,386 | 16,253 | 14,483 |
|  | Canu | 2.59/12.07/0.22 | 12.0 | 8.09 | 21,472 | 13,133 | 8,722 |
|  | NECAT | 2.85/1.33/2.14 | 9.01(11.35) | 45.85(25.67) | 23,600 | 15,944 | 7,158 |
| C. reinhardtii | raw reads | 14.84/--/-- | 15 | 1.16 | 54,409 | 46,812 | 4,231 |
|  | Canu | 4.61/59.40/0.078 | 5.35 | 76.05 | 53,891 | 45,934 | 726 |
|  | NECAT | 14.89/11.53/1.29 | 1.99(4.40) | 95.18(82.13) | 56,427 | 48,708 | 278 |
| O. sativa | raw reads | 63.40/--/-- | 15.6 | 0.49 | 56,325 | 50,847 | 24,205 |
|  | Canu | 15.23/43.20/0.35 | 7.99 | 44.42 | 55,010 | 49,612 | 4,413 |
|  | NECAT | 63.83/18.95/3.37 | 4.66(6.45) | 74.62 (51.49) | 56,573 | 51,141 | 3,511 |
| S. pennellii | raw reads | 132.74/--/-- | 18.49 | 1.7 | 24,801 | 22,226 | 127,808 |
|  | Canu | 37.53/88.8/0.42 | 9.69 | 34.04 | 21,653 | 19,364 | 5,511 |
|  | NECAT | 121.07/137.77/0.88 | 6.45 (9.23) | 63.04 (38.77) | 23,810 | 21,480 | 5,445 |

Size is the total number of base pairs in corrected reads. Time is the time of error correction, and the speed is the Size/Time. Error rate denotes the mean error rate of raw reads and corrected reads; <=5% denotes the percentage of reads with less than 5% error rate in total corrected read, values are the bracket are results of NECAT after the first correction; N50 and N75 are the length of read that reached the 50% and 75% of the total length of all reads; Read number with HERS denotes the number of reads that with at least one HERS (more than 50% error in the 500bp window). The reads that were used in evaluating the last three metrics (N50, N75 and Read number with HERS) of NECAT were corrected from longest 40x of raw dataset that were selected by Canu for correction by default, see Supplementary Note 6 for details.

**Table 2.**
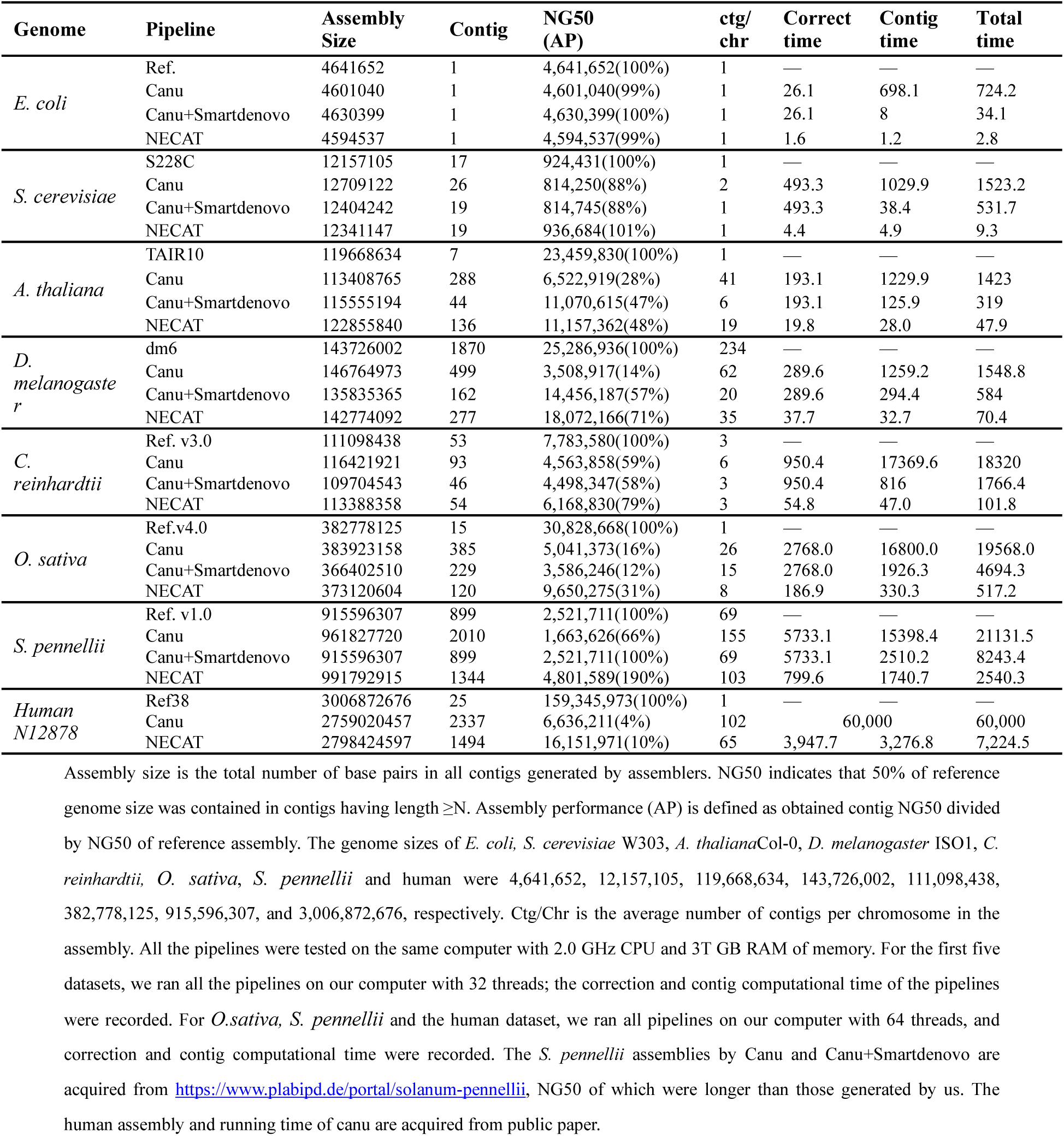
The quality and performance of long-read assembly with NECAT

**Table 3.** Performance of de novo assemblies before and after the bridging step of NECAT.

| Species | Stats | Count | Assembly Size | Max | Min | N25 | L25 | N50 | L50 | N75 | L75 |
| --- | --- | --- | --- | --- | --- | --- | --- | --- | --- | --- | --- |
| *E. coli* | Before | 1 | 4587234 | 4587234 | 4587234 | 4587234 | 1 | 4587234 | 1 | 4587234 | 1 |
|  | After | 1 | 4594537 | 4594537 | 4594537 | 4594537 | 1 | 4594537 | 1 | 4594537 | 1 |
| *S. cerevisiae* | Before | 20 | 12344710 | 1529545 | 37657 | 1087952 | 3 | 816246 | 6 | 581125 | 10 |
|  | After | 19 | 12341147 | 1529022 | 37657 | 1087471 | 3 | 936684 | 6 | 676549 | 10 |
| *A. thaliana* | Before | 150 | 122876764 | 14555777 | 4312 | 14075240 | 3 | 11149925 | 5 | 6575909 | 8 |
|  | After | 136 | 122855840 | 14566553 | 4312 | 14083693 | 3 | 11157362 | 5 | 7804579 | 8 |
| *D. melanogaster* | Before | 320 | 143000842 | 14922625 | 1303 | 12854107 | 3 | 9612127 | 6 | 2092117 | 14 |
|  | After | 277 | 142774092 | 21505040 | 1303 | 21396663 | 2 | 18072166 | 4 | 2925305 | 9 |
| *C. reinhardtii* | Before | 64 | 113293301 | 8997060 | 4161 | 6803426 | 4 | 5455837 | 9 | 3263676 | 16 |
|  | After | 54 | 113388358 | 9014332 | 4161 | 6812997 | 4 | 6168830 | 8 | 3374959 | 15 |
| *O. sativa Japonica Group* | Before | 167 | 372698321 | 22007406 | 3978 | 11903975 | 7 | 6099041 | 18 | 3370103 | 38 |
|  | After | 118 | 373827003 | 22086005 | 7816 | 13530842 | 6 | 10323607 | 14 | 5860244 | 25 |
| *S. pennellii* | Before | 1604 | 991874379 | 22857416 | 508 | 5921037 | 27 | 3465614 | 82 | 1668679 | 186 |
|  | After | 1344 | 991792915 | 22878582 | 508 | 6804220 | 22 | 4325703 | 67 | 2075284 | 151 |
| *Human* | Before | 2151 | 2791598215 | 50857421 | 500 | 26709700 | 19 | 15339800 | 55 | 7002196 | 124 |
|  | After | 1494 | 2798424597 | 73247802 | 500 | 31103549 | 15 | 16933776 | 47 | 8828295 | 102 |

Count is the total number of contigs in assembly. Assembly size is the total number of base pairs in assembly. N25/N50/N75 indicate that 25%/50%/75% of the assembly size is contained in the contigs of length $\geq N$ . The L25/L50/L75 are the number of contigs under the N25/N50/N75, respectively.

**Figure 1.**
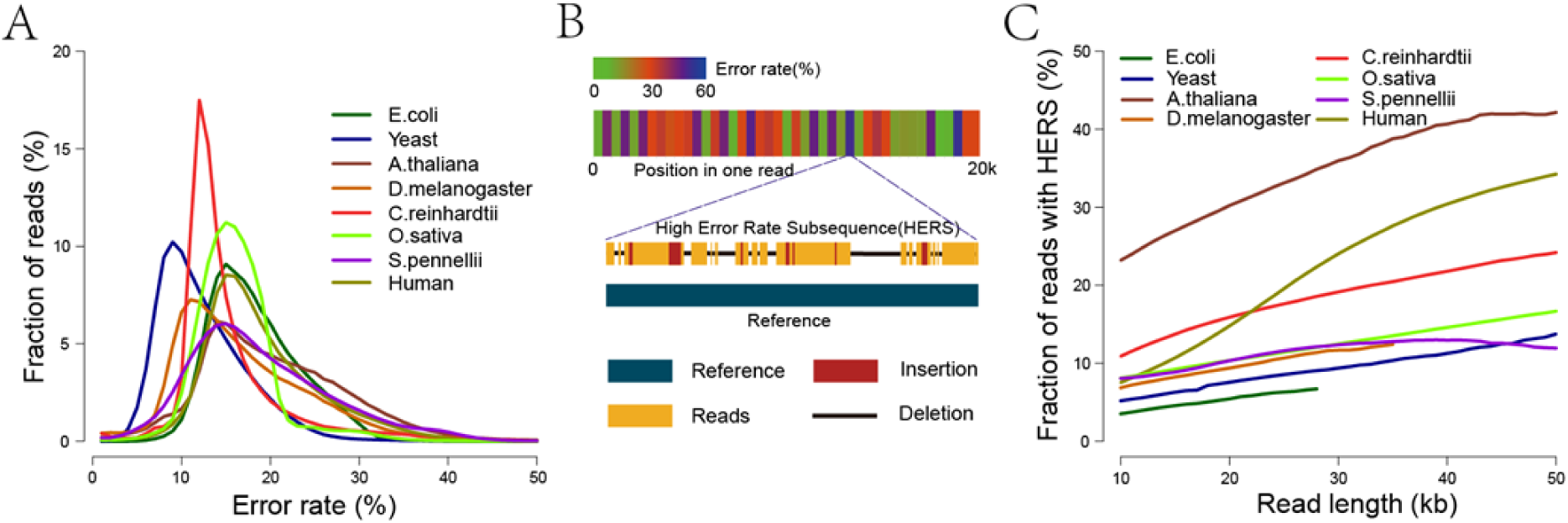
Error characteristics of eight Nanopore raw read datasets. (A) Error rate distribution of raw reads. (B) Error rates of subsequences in a Nanopore read (upper) and illustration of a high error subsequence in the read (bottom). (C) Plot of percentage of raw reads with high error rate subsequences (HERS, error rate more than 50% in 500 bp windows) against read length.

Next, we analyzed sequencing errors in each Nanopore read. We partitioned each read into 500-bp long subsequences and counted the error rate of each subsequence. Our results show that the error rates in each read are also broadly distributed (**Figure 1B**). Furthermore, on average, 3∼23% of raw reads longer than 10 kb have high error rate subsequences (HERS) with error rates greater than 50% (**Supplementary Table 3**). Overall, Nanopore reads produced by ultra-long library preparation techniques have a higher percentage of reads with HERS than those produced by normal library preparation techniques (23% vs. 3-11%). Additionally, the percentage of raw reads with HERS increased as read length increased (**Figure 1C**). Especially, in reads produced by ultra-long reads library preparation techniques, up to 45% of raw reads longer than 45 kb have HERS (**Figure 1C**). The HERS in Nanopore reads usually force the error correction tool to break long reads into shorter fragments, which eliminates the advantage of using long Nanopore reads for *de novo* assembly.

Furthermore, error rates of Nanopore reads sampled from different genome locations shared the same distribution except for those of *A. thaliana*, which showed slight variations among genome locations (**Supplemental Figure 1)**. These results indicate that Nanopore sequencing errors did not show genome-location bias. Therefore, a Nanopore dataset can contain both low and high error rate reads from the same location in a genome.

In summary, our analysis indicates that, unlike PacBio reads, Nanopore reads can contain HERS (especially in ultra-long raw reads), and show broad error rate distribution among reads and read subsequences.

### Adaptive selection of supporting reads for error correction

To correct a Nanopore read, we first collected supporting reads that overlap with it, then constructed the corrected read using a consensus of multiple-sequence-alignment of overlapped reads. An overlapping-error-rate threshold is usually set to select supporting reads. Due to broad distribution of sequencing-error rates among Nanopore reads, it is difficult to select supporting reads using a single global overlapping-error-rate threshold. Setting a low overlapping-error-rate threshold, such as 0.3 used for PacBio reads, does not generate enough supporting reads to correct Nanopore reads with high error rates (>20%); consequently, numerous Nanopore reads cannot be corrected. Conversely, setting a high overlapping-error-rate threshold (such as 0.6) to correct the majority of Nanopore reads results in markedly increasing of false supporting reads, which increases computational cost and reduce the accuracy of corrected reads. Furthermore, high overlapping-error-rate threshold can increase the number of high-error-rate supporting reads for low-error-rate template reads. This results in correcting low-error-rate template with high-error-rate supporting reads, which greatly reduces the accuracy of corrected low-error-rate reads.

To overcome the broad error-rate distribution of Nanopore reads, we used two overlapping-error-rate thresholds to select supporting reads after filtering via DDF scoring^11^ and k-mer chaining^19^ (**Online Methods**). First, we used a global overlapping-error-rate threshold to maintain the overall quality of supporting reads. Then, for each template read, we set an individual overlapping-error-rate threshold. The candidate reads were filtered if their alignment error rates were greater than either global or individual overlapping-error-rate thresholds. For low-error-template reads, the individual overlapping-error-rate threshold is less than the global threshold. Conversely, for high-error-rate template reads, the individual overlapping-error-rate threshold is greater than the global threshold. Using both global and individual overlapping-error-rate thresholds, we were able to maintain the quality of supporting reads for both low and high-error-rate template reads, thereby improving the accuracy of corrected reads. High-error-rate template reads that did not have enough supporting reads were discarded without correction.

### Progressive error correction of Nanopore reads

The supporting reads for error correction are selected according to average error rate of each template read. Since error rates for subsequences of each Nanopore read are also broadly distributed (**Figure 2A**), overlapping error rate between supporting reads and HERS can exceed the global threshold 0.5, which can affect the accuracy of corrected subsequences. Therefore, we developed a progressive method for correcting error prone Nanopore reads in two steps (**Online Methods**). We first corrected low-error-rate subsequences in a template read (**Figure 2B**). Then, we corrected high-error-rate subsequences (**Figure 2C**). In the first step, both corrected and uncorrected subsequences were outputted as a corrected read for the next step. After the first step, we corrected most Nanopore reads to high accuracy. This allowed us to obtain increased number of low-error supporting reads for high-error subsequences in the second step, thereby helping to correct high-error subsequences. After the second step, we outputted only the corrected subsequences. If a subsequence in a template read could not be corrected in the second step, it had either a high error rate or low coverage. Thus, one template read could be broken into multiple corrected reads.

**Figure 2.**
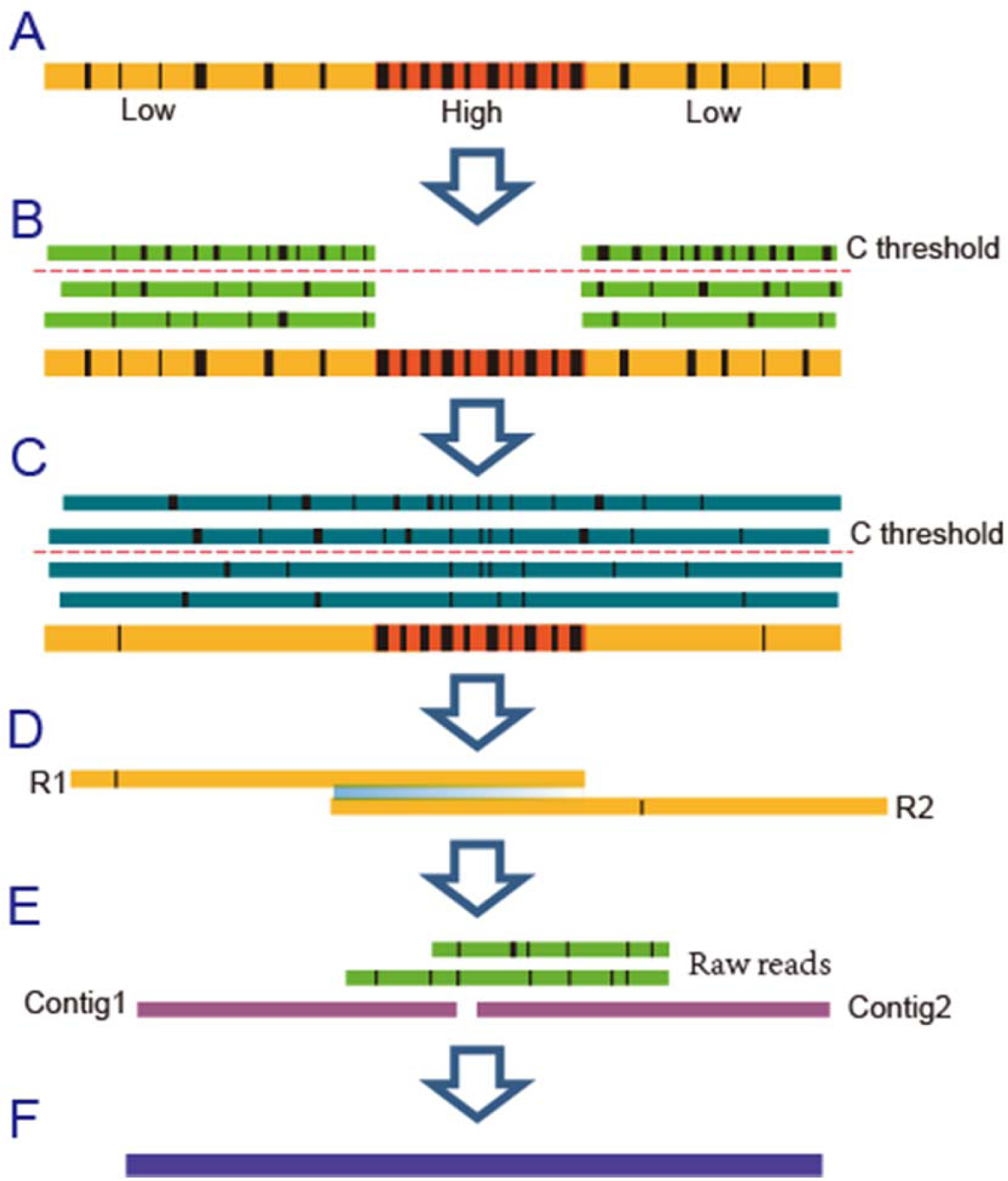
Illustration of progressive error correction and two-stage assembly methods of NECAT. (A) Input raw reads. (B) Error correction of low error rate subsequences. Only low error rate subsequences have supporting reads. (C) Error correction of high error rate subsequences. (D) Contig assembling using corrected reads. (E) Contig bridging using raw Nanopore reads. (F) Output final contigs.

Usually, twelve supporting reads are enough for error correction. Performing local alignments of supporting reads to template is computationally expensive, especially for long template reads. Although we selected 200 supporting reads for each template read, it is unnecessary to align all these supporting reads when there are enough reads available for error correction. Thus, we used a coverage count array (CCA) to record the number of supporting reads that covered each base of the template read. For template read covered by a sufficient number of support reads, we did not perform local alignment of supporting reads to this region anymore (**Online Methods)**.

### Progressive assembly of Nanopore reads

The long length of Nanopore reads is a significant advantage for *de novo* genome assembly. However, HERS inside long Nanopore reads usually fail to be corrected, leading to the splitting of long Nanopore reads into several shorter corrected reads. Using only corrected reads for genome assembly abolishes the advantage presented by the long length of Nanopore reads. In this study, we developed a two-step progressive genome assembler for Nanopore reads. In the first step, we generated high quality contigs using corrected reads (**Figure 2D**). In the second step, we bridged the contigs using original Nanopore reads to generate final scaffolds (**Figure 2E**). The lost contiguity in contigs, caused by HERSs in raw reads, is thereby filled in the second step of the process. Therefore, genome contiguity is improved by maximizing the usage of all raw reads. Our two-step assembly process is similar to process using SMS reads for scaffolding^20^.

Meanwhile, even after error correction, sequencing error rates of corrected Nanopore reads (1.5-9%) are still higher than those of corrected PacBio reads (less than 1%). Moreover, the error rates of corrected reads also show a relatively broad distribution (**Supplementary Note 6 and Supplementary Table 4**). To obtain high quality contigs, we needed to select high-quality overlaps between corrected reads because low-quality overlaps increase the difficulty of assembly and introduce errors into assembly results. Similar to the process used for selecting supporting reads for error correction, we employed both global and individual thresholds to overcome the broad-error-rate distribution for the filtering of low-quality overlaps (**Online Methods)**.

### Performance of NECAT error correction

We assessed the performance of NECAT error correction using Nanopore raw reads of seven species: *E.coli, S. cerevisiae, D. melanogaster, A. thaliana, C. reinhardtii, O. sativa*, and *S. pennellii* with respect to correction speed, corrected data size, accuracy and continuity of corrected reads, as well as the number of reads with HERS in corrected reads (**Supplementary Note 6**). As shown in **Table 1**, NECAT correction speeds were 2.1-16.5 times faster than those of Canu for Nanopore reads of these seven species. The sizes of corrected reads for *E.coli, S. cerevisiae, D. melanogaster, A. thaliana, C. reinhardtii, O. sativa*, and *S. pennellii* were 102.2%, 83.4%, 90.6%, 92.5%, 100.3%, 100.7% and 91.2% of their raw reads, respectively, while Canu only corrected the longest 40X raw reads and obtained 15.9%, 39.8%, 57.7%, 84.1%, 31.1%, 24.0%, and 28.3% corrected reads from their raw reads, respectively.

NECAT was able to obtain high-accuracy corrected reads. After the first step, average error rates for *E.coli, S. cerevisiae, D. melanogaster, A. thaliana, C. reinhardtii, O. sativa*, and *S. pennellii* datasets were 4.27%, 3.08%, 7.03%, 11.35%, 4.40%, 6.45%, and 9.23% respectively; these were less than the average error rates of reads corrected by Canu, which were 7.06%, 3.13%, 8.15%, 12.05, 5.35%, 7.99%, and 9.69% respectively. After the second step, average error rates for seven datasets were further reduced to 2.23%, 1.53%, 4.89%, 9.01%, 1.99%, 4.66%, and 6.45%, respectively.

The maximum overlapping error rate between corrected reads is usually set to 10% during assembly. Thus, the higher the percentage of corrected reads having less than 5% error, the more reads can be used for assembly. As shown in **Table 1**, the percentages of NECAT’s corrected reads having error rate less than 5% error for seven data sets were 99.34%, 95.04%, 72.03%, 45.85%, 95.18%, 74.62%, and 63.04% respectively, which were significantly higher than those of reads corrected by Canu.

The progressive correction strategy in NECAT also allowed us to correct more HERS and maintain the contiguity of reads. N50s for NECAT-corrected reads of the seven datasets were 105.1%, 90.5%, 98.0%, 100.9%, 103.7%, 100.4%, and 96.3%, respectively, of N50s for their corresponding raw reads, indicating that NECAT could preserve the contiguity of raw reads. Conversely, N50s for the reads corrected by Canu were 91.9%, 30.4%, 85.8%, 91.8%, 99.0%, 97.7% and 87.3% of the corresponding raw reads, which was less than those of NECAT-corrected reads. Another evidence that progressive correction strategy in NECAT can improve the correction of HERS is that the number of reads with HERS has been reduced. After two-step correction using NECAT, the numbers of reads containing HERS in the seven corrected datasets were 1, 268, 3,481, 7,158, 278, 3,511, and 5,445 respectively, while Canu-corrected datasets had 1, 4,820, 6,523, 8,722, 726, 4,413 and 5,511 reads containing HERS. These results indicate that NECAT outperformed Canu in correcting sequencing errors in Nanopore raw reads.

### Performance of NECAT *de novo* assembler

We compared NECAT to two widely used correct-then-assemble pipelines, Canu and Canu+smartdenovo, for *de novo* assembly of Nanopore reads (**Supplementary Note 7)**. We assembled genomes of *E. coli, S. cerevisiae, A. thaliana, D. melanogaster, C. reinhardtii, O. sativa* and *S. pennellii* using the longest 40X reads of each dataset, and assembled 35X Nanopore data for the human NA12878 genome using NECAT only. As shown in **Table 2**, NECAT was 8.3-258.2 times faster than Canu, while showing 8.8-577.5 times speedup during the assembly step. Canu employs a high overlapping threshold (14.4%) in its overlapIncore tool for Nanopore reads (a low threshold of 6% is used for assembling PacBio reads), which may greatly increase the time cost of local alignments. The Canu+smartdenovo pipeline replaces the assembly step of Canu with smartdenovo, which significantly reduces running time. NECAT was still 3.2-57.0 times faster than Canu+smartdenovo on seven datasets. The high accuracy of corrected reads outputted by NECAT allowed us to use a more rapid overlapping approach.

We then assessed the quality of assembled contigs with respect to assembly size, NG50, number of contigs, and average number of contigs > 200 bps per chromosome (ctg/chr). For *E. coli*, all three pipelines recovered the complete genome in just one contig. For *S. cerevisiae*, NECAT outperformed Canu and Canu+smartdenovo with 101% assembly performance and a near perfect contiguity with only 19 contigs. For *A. thaliana*, NECAT reported 136 contigs and an NG50 of 48% assembly performance, which was similar to that of Canu+smartdenovo (47% assembly performance) and markedly better than that of Canu (28% assembly performance). For *D. melanogaster*, NECAT reported 277 contigs and obtained the best NG50 performance (71% assembly performance) compared with those of Canu (14% assembly performance) and Canu+smartdenovo (57% assembly performance). For *C. reinhardtii*, NECAT reported 54 contigs and the best NG50 performance (79% assembly performance). For *O. sativa*, NECAT reported 120 contigs and the best NG50 performance (31% assembly performance), which was markedly better than those of Canu (16% assembly performance) and Canu+smartdenovo (12% assembly performance). For *S. pennellii*, NECAT reported 1344 contigs and the best NG50 performance (190% assembly performance), which was 1.90 and 2.88 times greater than those of Canu+smartdenovo (100% assembly performance) and Canu (66% assembly performance). For human NA12878, NECAT report 1494 contigs and 16.93 Mbp NG50 (30% assembly performance), which was 2.43 times longer than that reported by Canu. Furthermore, NECAT assembled the human NA12878 genome in only 4.7 days on a single 64-threaded computer.

We next assessed the effect of contig-bridging in NECAT assembly. As shown in **Table 3**, the number of contigs was significantly reduced in the assembly of *A. thaliana, D. melanogaster, C. reinhardtii, O. sativa, S. pennellii* and *H. sapiens* genomes after contig-bridging of raw reads. For *D. melanogaster* and *S. pennellii* contig-bridging also significantly increased the N50 of assembly. These results indicate that contig-bridging can significantly improve the contiguity of assembly.

We further compared NECAT assembler with widely used assemble-then-correct assemblers: Miniasm, Smartdenovo, Wtdbg2, and Flye (**Supplementary Text 1 and Note 7)**. NECAT has similar time costs as those assemble-then-correct assemblers, but obtains better assembly results, especially for complex genomes (**Supplementary Text 1)**. We also validated our assemblies by comparing them to reference genomes. The quality of NECAT-generated assemblies were comparable to those of the other correct-then-assemble pipelines and better than assemble-then-correct assemblers (**Supplementary Text 2**).

### De novo genome assembly of retinoblastoma cell line WERI

To further evaluate the performance of NECAT in large-genome assembly, we sequenced a cell line called WERI, which is derived from human retinoblastoma^21^. We generated 210 Gb (82 folds) of raw reads from three flowcells using Nanopore PromethION. The WERI genome assembled by NECAT has an N50 of 29M. To the best of our knowledge, this is the best N50 value for the assembly of human genome using the general library of the Nanopore sequencing platform.

We aligned the WERI assembly to human reference genome hg38 using MUMmer (v4.0)^22^. The dotplot figure shows that the WERI assembly is structurally consistent with reference genome except for minor structural variations (**Supplementary Note 8** and **Supplementary Figure 2**) and the tiling figure shows the continuity of the assembly (**Figure 3**). We also used bowtie2^23^ to align an Illumina dataset for the WERI cell line onto a WERI assembly and hg38 human reference genome. The mapping rate of the WERI assembly (99.1%) was better than that of hg38 human reference genome (98.0%).

**Figure 3.**
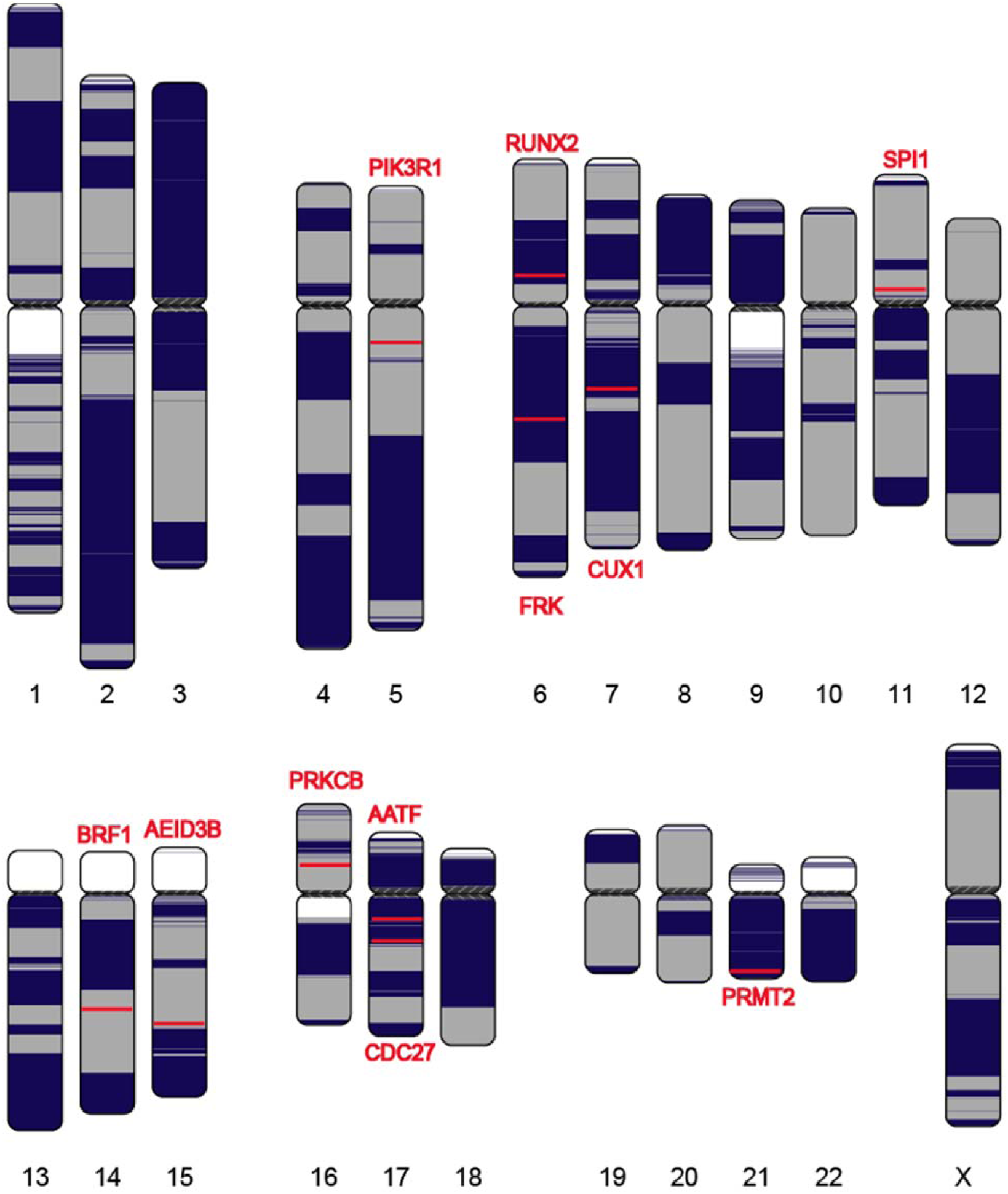
Continuity analysis of the assembly of WERI cell line using Nanopore reads. Human chromosomes are painted with assembled contigs using the ColoredChromosomes package. Alternating shades indicate adjacent contigs (each vertical transition from gray to black represents a contig boundary or alignment breakpoint).

We then identified and validated structural variants (SVs) in the WERI assembly. We detected 11,725 SVs (≥10 bp) in the WERI assembly by aligning it to hg38 human reference genome using Nummer (v4.0). We also detected SVs from raw Nanopore long reads and Illumina short reads for the WERI cell line using Sniffles^24^ and LUMPY^25^, respectively (**Supplementary Note 8**). 7210 SVs are commonly detected using WERI assembly and raw Nanopore reads, while only 1117 SVs are commonly detected using WERI assembly and NGS (**Supplementary Figure 3 and Supplementary Table 5**). Furthermore, 90% of unique small SVs (<1000 bp) detected using Nanopore raw reads were able to be found in the WERI assembly, indicating that the assembly can reduce false positives for small SVs (<1000 bp) (**Supplementary Table 5**).

Next, we examined genes associated with the identified SVs. We found 2843 annotated genes associated with 7210 SVs identified using both WERI assembly and raw Nanopore reads. 209 of 2843 genes are reported in Phenolyzer^26^ and are associated with retinoblastoma (**Supplementary Table 6**). Among 66 genes, the gene *PRKCB*, which is scored as high as 0.8901 in Phenolyzer^26^, was reported to be involved in retinoblastoma protein phosphorylation^27^. Among the 209 genes, there are eight genes (*AATF*, PRKCB, *PRMT2, FRK, PIK3R1*, CUX1, RAC2, IGF1) with a Phenolyzer score greater than 0.5, and six of eight genes are associated with retinoblastoma as reported in PubMed. These results indicate that NECAT can provide high quality assembly for reliable identification of SVs.

## Discussion

Currently, applying Nanopore reads in genomic studies is difficult because of the complex errors within these reads. In this study, our analyses have shown that Nanopore reads contain high-error rate subsequences, and errors are broadly distributed among Nanopore reads and in subsequences of a read. This broad error distribution complicates selection of supporting reads during the error-correcting process. In traditional error-correction methods, the threshold used to select supporting reads can be set too strict or too lenient; the former cannot select enough supporting reads for correction, while the latter generates too many low-quality reads that affect the accuracy of corrected reads. Furthermore, traditional error correction methods cannot correct the high-error-subsequences in Nanopore reads and generally break Nanopore reads into multiple short corrected reads.

In this study, we developed NECAT, which includes novel methods such as progressive error correction, adaptive supporting reads and alignment selection, and two-stage assembly, to overcome the errors characteristic of Nanopore reads. The novel error-correction tool in NECAT, which is 2.1-16.5 times faster than that of Canu, can correct Nanopore reads to high accuracy, while maintaining the contiguity of Nanopore reads. The novel assembly tool in NECAT is at least 1.4 times faster than other assembly pipelines with enhanced or comparable assembly performance. The high performance shown by NECAT suggests that the high error rate of Nanopore reads can be overcome by the development of new algorithms with respect to error characteristics.

Structural variations identified via raw Nanopore reads usually have a high false-positive rate. Here, we show that these false positives can be reduced considerably by using a high-quality assembly of Nanopore reads for detection of structure variation. Our results show that NECAT is a useful tool for error correction and assembly of Nanopore reads, and for detection of structure variation.

## Data sources

We used nine datasets to evaluate the performance of NECAT. Among these datasets, those for *Saccharomyces cerevisiae, Oryza sativa and Homo sapiens* (the WERI human retinoblastoma cell line) were generated using our in-house sequencing, while the other four were obtained from public websites. The details on the data used in this study are reported in **Supplementary Notes 1-4**.

## Accession codes

All processed files for assembly and analysis code used in this study are available from http://www.tgsbioinformatics.com/necat. All source codes for NECAT are available from https://github.com/xiaochuanle/NECAT.

## ACKNOWLEDGMENTS

We thank all those who generated and freely released the data analyzed in our present study. This study was funded in part by the National Natural Science Foundation of China (grant numbers 31871326, 31701146, 91953122, 6832019, 61420106009, 81530028, 81721003). We thank the Local Innovative and Research Teams Project of Guangdong Pearl River Talents Program, Clinical Innovation Research Program of Guangzhou Regenerative Medicine and Health Guangdong Laboratory (grant number 2018GZR0201001); the State Key Laboratory of Ophthalmology, Zhongshan Ophthalmic Center, Sun Yat-sen University. This work was supported in part by the U. S. National Institute of Food and Agriculture (NIFA; grant number 2017-70016-26051) and U.S. National Science Foundation (NSF; grant number ABI-1759856) to F. L.

## AUTHOR CONTRIBUTIONS

C.L.X., Y.Z.L., J.X.W., and F.L. conceived and designed this project. Y.C. and C.L.X. conceived, designed, and implemented the consensus algorithm. F.N. and C.L.X conceived, designed, and implemented the progressive assembly algorithm. F.N. and Y.C. integrated all the programs into the NECAT pipeline and provided documentation. S.Q.X., C.L.X., Y.X.W., J.F.X. and Q.D. ran analyzed genome assemblies and analyzed the performance of algorithms developed in this study. T.B., Z.J.H., D.P.W. and L.J.H. coordinated data release and assisted with executing the pipeline. F. L., Y.C., and F.N. performed theoretical analysis of the algorithms developed in this study. F.L., C.Y., F.N., S.Q.X., Y.F.Z. and C.L.X. wrote the manuscript. All authors have read and approved the final version of this manuscript.

## COMPETING FINANCIAL INTERESTS

The authors have no competing financial interests to declare.

## ONLINE METHODS

### The architecture of NECAT

The NECAT pipeline was designed as a high-performance assembler for Nanopore reads. To overcome the high-error-rate of Nanopore reads, we developed several novel methods, including progressive error correction, adaptive supporting reads and alignment selection, and two-step assembly. The NECAT pipeline contains four modules (Supplementary Figure 4): preprocessing, correction, trimming, and assembly. The preprocessing module filters short and ill-formed reads. The correction module uses a progressive strategy to correct Nanopore reads in two steps. The trimming module removes low-quality subsequences from corrected reads. The assembly module builds a string graph to assemble the genome in two steps. These four modules can be run in series to finish assembly, or can be operated independently. Currently, NECAT is the most efficient assembler for large genomes from Nanopore reads. NECAT also significantly improved the contiguity of the assembled genome.

### Progressive error correction of Nanopore reads

The broad distribution of sequencing-error-rates among Nanopore reads, and within a single Nanopore raw read, is the reason for why traditional iterative error-correction methods usually fail with Nanopore data. In this study, we develop a novel method for correcting Nanopore reads. Our progressive error correction method involves two steps. First, we correct the low-error-rate subsequences (LERS) in a read. Then, we correct the high-error-rate subsequences (HERS) in that read using a more sensitive approach. Both steps include the same four sub-steps: i) selection of candidate reads, ii) determination of alignment-quality threshold, iii) selection of matched reads, and iv) correction of the read. The sub-steps i, ii, and iv are the same for both steps. We use different methods to select matched reads for each template to be corrected in sub-step iii of the two steps. In first step, we use a strict selection method to choose matched reads for the low-error-rate portions of template read. In second step, we use a lenient method to choose matched reads for the high-error-rate portions of template read.

### Selection of candidate reads

For each read to be corrected, we select candidate reads that have overlap with that read. For each pair of reads, we first use the distance difference factor (DDF^11^) to select a seed k-mer pair with the highest score, which serves as a reliable start position for local alignment. However, the wide distribution of error rates decreases the sensitivity of the DDF score for two k-mer pairs that are far apart; this may introduce false positives (**Supplementary Figure 5A**). To remove false positives, we gather all k-mer pairs that support the seed k-pair during DDF scoring. We sort all k-mer pairs, including the seed k-mer pair, with respect to their positions and then chain them together^19^. The chaining process examines the relative positions of k-mer pairs and helps to filter out false positives (**Supplementary Figure 5B**). We then update the DDF score of the seed k-mer pair with remaining k-mer pairs, which further improves the sensitivity of candidate selection. We record the positions of the first and last k-mer pairs in the chain as the approximate mapped positions of candidate read. These two positions, together with the DDF score of the seed k-mer pair, are used for further filtering of redundant candidates and identifying HERS.

### Determination of individual alignment-quality threshold for each template read

We select high-quality supporting reads that are used for the correction of each template read. However, broad error rate distribution makes it difficult to use a single global threshold for selection of supporting reads. Besides setting a global overlapping-error-rate threshold to 0.5, we also compute a local individual overlapping-error-rate threshold for each template read. For each template read, we use 50 candidate reads with top DDF scores for local alignments. If a local alignment contains more than 60% of template or candidate read length, we record the alignment, and the difference between template and candidate read. If we have *n*(0 ≤ *n* ≤ 50) recorded alignments and their differences are *d*_1_, *d*_2_,…, *d*_*n*_, We compute their average difference 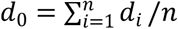 and standard deviation 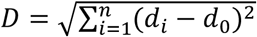. Then, we set the alignment quality threshold as *d* = *d*_0_ − 5*D*. This threshold provides a lower alignment quality bound for low error template reads.

### Selection of matched reads

For each read template, we select 200 candidate reads with top DDF scores for local alignment. We use different alignment methods in first and second steps. In the first step, we use blockwise alignment algorithm for aligning supporting reads to the template read. We perform local alignment from the seed k-mer pair in both directions. Thus, we first obtain two semi-global alignments, and then the two alignments are merged into one. Starting from the seed k-mer pair, we partition both template and candidate reads into equal-sized blocks 500 bp in length. We then use the Edlib algorithm^28^ to successively align each pair of blocks. The aligning process terminates if the alignment error between a pair of blocks is greater than 50%, or if the alignment algorithm reaches the end of a template or candidate read. Because blockwise alignment terminates when either block from template or candidate has a high error rate, we can only obtain alignment between low-error-rate subsequences in this step.

In the second step, we use multiple alignment methods to obtain long alignments between templates and candidate reads. We first use the blockwise approach to align candidate reads to a template. If blockwise alignment terminates early due to presence of a high-error-rate region inside the template or candidate read, we use the DALIGN algorithm^29^ to re-align the candidate read to template. However, alignments produced via DALIGN, running with a large difference threshold of 0.5, are usually too coarse. To refine the alignment result of DALIGN, we then use the Edlib algorithm to perform a global alignment on the mapped subsequences output by DALIGN to get a more correct alignment.

Performing a local alignment of supporting reads to template is computationally expensive, especially for long-template reads. Usually only dozens of alignments are enough for error correction. Thus, it is unnecessary to align all 200 candidate reads if we have enough supporting reads for error correction. Here, we use a coverage count array (CCA), which is an integer array possessing the same length as that of template read, to record the number of candidate reads that cover each base of the template read. Prior to aligning a candidate read to the template read, we examine the values of CCA elements between the mapped positions for the approximate start and end of candidate read on a template. If all these values are greater than a user set threshold *C*, we would know that the corresponding region in template read has been covered by enough candidate reads and there is no need to perform the local alignment of this candidate read. If the alignment difference is less than the alignment quality threshold *d*, we would increase every value of CCA between the start and end template mapped positions by one. We use a default value of 12 for threshold *C*.

### Correction of Nanopore reads

After selecting matched candidate reads, we use the FALCON-sense consensus algorithm^9^ to correct each subsequence of the template read that is covered by enough candidate reads. In the first step, we replace these subsequences with corrected subsequences. Then, we output the whole template, including corrected subsequences and uncorrected subsequences, as a corrected read for the next step. HERS are corrected in the next step. In the second step, we only output corrected subsequences, meaning that one template may produce more than one corrected read. If a subsequence in a template read cannot be corrected in the second step, it either has too high of an error rate or low coverage.

### Trimming of low-quality subsequences

Long Nanopore reads may still contain HERS even after error correction, which can greatly affect the quality of assembly. Thus, low-quality subsequences need to be trimmed before assembly. We only select 40X coverage longest corrected reads for trimming and future assembly. First, we perform pairwise alignment on selected Nanopore reads using the trimming module of MECAT^11^. Because even corrected Nanopore reads may have a relatively high error rate, we use the sensitive DALIGN algorithm to replace the original diff algorithm in the MECAT trimming module before performing local alignments. After pairwise alignment, we gather high-quality overlaps with more than 90% identity for each read. If every residue of a read is covered by at least one overlap, the read is designated as a complete read. On the other hand, if there are subsequences without overlap coverage in a read, we trim it to its longest covered subsequence, which is called a trimmed read.

After trimming, the reads are usually subjected to another pairwise alignment. Our experiments show that less than 10% of corrected reads are trimmed, therefore, it is unnecessary to pairwise align 90% of untrimmed reads. Thus, we store complete reads and trimmed reads separately after trimming. Pairwise alignments are only performed between complete reads and trimmed reads, and between trimmed reads. The results of these pairwise alignment, together with complete reads, trimmed reads, and results of original pairwise alignments between complete reads, are fed into the assembly module.

### *De novo* assembly of Nanopore reads

Although the long length of Nanopore reads helps improve genome assembly, the relatively high error rate of these reads renders genome assembly difficult. Here, we developed a new assembly tool that is particularly useful for Nanopore reads because it can overcome the high error rate of these reads. Our assembly module in NECAT consists of three steps: filtering of low-quality read overlaps, contig assembly, and contig bridging. We use multiple quality-control measures to filter out low-quality overlaps between Nanopore reads. Then, we construct a directed string graph and solve the graph to generate contigs. Finally, we bridge the contigs using original reads to generate the final scaffolds.

### Filtering of low-quality read overlaps

Low-quality overlaps complicate assembly and introduce errors into assembly results. In NECAT, we use multiple thresholds to control the identity, overhang, and coverage of overlaps in order to filter out low-quality overlaps. For each read, we determine the coverage of each base according to its overlaps. Then, we calculate the minimum coverage (*c*_*min*_), maximum coverage (*c*_*max*_) of bases, as well as the difference between minimum coverage and maximum coverage (*c*_*diff*_). If its *c*_*min*_ is less than predefined threshold, *min_coverage*, or *c*_*max*_ is larger than predefined threshold, *max_coverage*, or *c*_*diff*_ is larger than predefined threshold, *max_diff_*coverage, the read and its overlaps are removed. The details on coverage threshold settings are provided in **Supplementary Note 9**. Because of broad error distribution among different reads, we use both global and local threshold, instead of a single global threshold, for quality control of overlap identity and overhang. For a high-quality read, the average quality of its overlaps needs to be higher than global average; therefore, we set the local threshold to filter out overlaps having relatively low quality. For a low-quality read, the average quality of its overlaps needs to be lower than global average; we then use the global threshold to filter out low-quality overlaps for that read. This strategy allows us to filter out overlaps with relatively low quality for each read, and to maintain the overall quality of all the overlaps. Details on setting global and local thresholds for overlap identity and overhang are provided in **Supplemental Note 9**.

### Contig assembly

Next, we construct a directed string graph and remove transitive edges using Mayer’s algorithm^30^. We mark the best out-edge and the best in-edge of each node based on overlap lengths of the edges. The edges that are not marked as best out-edge or best in-edge are removed^31^. We also remove ambiguous edges (tips, bubbles, and spurious links) in the graph. We then identify linear paths from the graph and generate contigs. When there is a branch, we break the path to generate multiple contigs. This strategy can reduce the possibility of mis-assembly.

### Contig bridging

During error correction, long reads with high-error subsequences are cut into multiple shorter reads, which eventually leads to discontinuity of contigs. It is possible to relink contigs using long raw reads^20^. First, we align the long raw reads to contigs. Two contigs may have an overlap that is of low quality; this overlap is filtered before construction of a string graph. A raw read can either fill the gap between two contigs, which is then called a gap read, or overlap with the overlap of two contigs, which is then called an overlap read. For each raw read, we record the gap or overlap length between the mapped positions on the ends of the two contigs. For each pair of contigs, the raw reads connecting them are grouped as those connecting in same orientation or those connecting in different orientations. In each orientation group, we cluster the raw reads based on their gap/overlap lengths. If the difference between the gap/overlap lengths of two raw reads is less than threshold (default value is 1000 bp), we assign them into same cluster. And we assigned a score to each raw read, which is the sum of the products of identity and length of overlaps between the raw read and the pair of contigs. The read cluster with the largest sum of scores is chosen as the link for the contig pair.

After identifying links between contig pairs, we create a string graph in which contigs are nodes and links between the contigs are edges. The weight of each edge is set to the link score. We simplify the graph again by removing transitive edges. Then, we traverse the graph and identify linear paths as final contigs. A raw read from the link is selected to fill the gap between contigs.

### Error distribution analysis

We analyzed error distribution in Nanopore datasets for *E. coli, S. cerevisiae, A. thaliana, D.* melanogaster, *C. reinhardtii, O. sativa* and *S. pennellii*. Our results indicate that the sequencing error rate of Nanopore reads was high at 10-30%, which helped us refine our algorithm for the NECAT platform and provide insights into why the existing correction algorithms are not suitable for the correction of Nanopore reads. Details are provided in **Supplementary Note 5**.

### Evaluation

We compared our error correction tool with those provided in Canu. We also systematically evaluated the assembly tools provided in NECAT by comparing them with those of Canu and Canu+smartdenovo. Details of these comparisons are reported in **Supplementary Notes 6-7**,**10**.

