## Supplementary Notes for "Fast and accurate assembly of Nanopore reads via progressive error correction and adaptive read selection"

### Supplementary Text 1：Comparison with assemble-then-correct assemblers

We compared our NECAT assembler with widely used assemble-then-correct assemblers: miniasm[1], smartdenovo, wtdbg2[2], and Flye[3] (**Supplementary Note 7)** using Nanopore data of *E. coli*, *S. cerevisiae*, *A. thaliana*, *D. melanogaster*, *C. reinhardtii*, *O. sativa* and *S. pennellii*. In general, assemble-then-correct assemblers run fast but obtain relatively poor assembly results.

As shown in **Supplementary Table 7**, running time of NECAT was similar to those of assemble-then-correct assemblers. wtdbg2 was the fastest assembler on all datasets except on *A. thaliana*. NECAT was 1.6-14.2 times faster than Smartdenovo on first six datasets. NECAT was 1.2-1.8 times faster than Flye on datasets of *A. thaliana*, *D. melanogaster*, *C. reinhardtii*, *O. sativa* and *S. pennellii* and was 224.8 and 22.1 times faster than Flye on datasets of *E. coli* and *S. cerevisiae*. NECAT was 1.2-6.8 times faster than miniasm on datasets *S. cerevisiae*, *C. reinhardtii* and *O. sativa*, but miniasm was 8.1 and 1.9 times faster than NECAT on datasets *A. thaliana* and *D. melanogaster.*

We then assessed the quality of assembled contigs from four aspects: assembly size, NG50, number of contigs and the average number of contigs > 200 bps per chromosome (ctg/chr). For *E. coli*, all assemblers recovered the complete genome in just one contig. For *S. cerevisiae*, NECAT reported 19 contigs and an NG50 of 101% assembly performance, which is similar to those of Flye (102% assembly performance) and Smartdenovo (101% assembly performance). For *A. thaliana*, NECAT reported 136 contigs and an NG50 of 48% assembly performance, which is similar to those of

Flye (51% assembly performance) and miniasm (48% assembly performance). For *D. melanogaster*, NECAT reported 277 contigs and obtained the best NG50 performance (71% assembly performance), while Flye has the second-best assembly performance (47% assembly performance). For *C. reinhardtii,* NECAT reported 54 contigs and the second best NG50 performance (79% assembly performance), closing to the best NG50 performance Flye (84% assembly performance). For *O. sativa*, NECAT and miniasm outperformed other assemblers with 31% assembly performance. For *S. pennellii*, NECAT reported 1344 contigs and 4.80Mbp NG50, which is 2.4 times longer than the second best NG50 reported by Flye (1.97Mbp). With the high accuracy of corrected reads output by NECAT, NECAT can take advantage of the correct-then-assemble strategy and performs better in assembling complex genomes.

**Supplementary Text 2：Validating assemblies from Nanopore reads**

We further validated our assemblies by comparing them to reference genomes. The assemblies of the first five genomes were polished by nanopolish[4] and pilon[5] and The assemblies of O. sativa and Human genomes were polished by Racon[6] (**Supplementary Note 10)**. First, we mapped the assemblies of *E. coli, S. cerevisiae, A. thaliana, C. reinhardtii*, *D. melanogaster*, *O. sativa*, and Human N12878 from Nanopore reads to corresponding reference genomes using MUMmer (v4.0)[7], then evaluated the mapping results using GAGE scripts[8]. Except for the presence of minor structural variations, most assemblies were structurally consistent with reference genomes (**Supplementary Figures 6-12**). Most assemblies are good collinearity with reference genomes, except the assemblies of *A. thaliana* and *D. melanogaster* generated by wtdbg2, *C. reinhardtii* generated by Canu+smartdenove and smartdenove. Second, for S. pennellii[9], we mapped the assembly of NECAT to the assemblies of the other pipelines from public paper using MUMmer (v4.0)[7], our assembly were structurally consistent with the assemblies except for the presence of minor structural variations (**Supplementary Figure 13**), since NG50 of NECAT-generated assembly was much longer than the original reference genome that was generated by Canu+Smartdenvo[9]. The tiling figure also shows that continuity of human N12878 assembly generated by NECAT was better than that generated by Canu (**Supplementary Figure 14**).

**Supplementary Table 8** provides GAGE[8] accuracy metrics for the assemblies of *E. coli, S. cerevisiae, A. thaliana, C. reinhardtii*, and *D. melanogaster*. The numbers of single-nucleotide polymorphisms (SNPs) and large indels (>10bps) in the genomes assembled by Canu, Canu+Smartdenovo, Smartdenovo, miniasm, wtdbg2, Flye, and NECAT were similar. Assemblies reported by NECAT maintained at least 99.30% coverage of their reference genomes.

We then mapped 17,294 annotated genes from *D. melanogaster*[10, 11] onto its three assemblies (**Supplementary Note 11**). A total of 16,402, 16,438, 16,368, 16,507, 16458, 16,396 and 16,412 genes were mapped onto a single contig of assemblies generated using Canu, Canu+smartdenovo, Smartdenovo, miniasm, wtdbg2, Flye and NECAT in a single alignment; 15,926, 15,956, 15,979, 16,017, 16,084, 16,075 and 16,053 of these genes showed over 99% identity. This indicates that the quality of NECAT assembly was comparable to those of the other pipelines.

Solving repeat regions is the most important task in genome assembly. We first evaluated three assemblies of *D. melanogaster* by comparing the completeness of transposable element (TE) families[12] (**Supplementary Note 11**). Of the 5,433 annotated TEs from FlyBase, NECAT assembly contained 5,304 TEs, in which 4,001 were aligned perfectly to the reference genome. Flye and wtdbg2 assemblies contained only 3840 and 3831 TEs aligned perfectly to the reference genome, which were less than other assemblies. We then examined two TE families: *roo* and *juan*. Using NECAT assembly, we aligned 134 of the 138 copies in the *roo* family, of which 118 were aligned perfectly. The 11 elements of *juan* family were also aligned perfectly. These results are similar to those obtained using other pipelines except miniasm (**Supplementary Table 9**). miniasm assembly only contained 77 perfectly aligned elements of *roo* family and 9 perfectly aligned elements of *juan* family.

We also examined telomeric repeats of 16 chromosomes in NECAT assembly of *S. cerevisiae* (**Supplementary Note 12**). We mapped 14 out 16 telomeric repeats to both ends of each chromosome. One telomeric repeat was mapped onto two chromosomes, and the other telomeric repeat was mapped to one end of a chromosome. Our results were similar to those obtained using assemblies generated by other pipelines except wtdbg2. wtdbg2 assembly contained 8 telomeric repeats mapped onto two chromosomes and 5 telomeric repeats mapped to one end of a chromosome (**Supplementary Table 10**). Both TE of *D. melanogaster* and telomeric repeat of *S. cerevisiae* analyses demonstrated that NECAT could accurately reconstruct repeat sequences.

**Supplementary Note 1: Cell culture and sequencing materials**

Datasets for eight species (*E. coli*, *S. cerevisiae*, *A. thaliana*, *D. melanogaster*, *C. reinhardtii*, *O. sativa*, *S. pennellii* and *H. sapiens*) were used to train and test our algorithm. Among these, four datasets (*S. cerevisiae*, *C. reinhardtii*, *O. sativa Japonica Group*, and retinoblastoma cell line WERI) were cultured and sequenced using MinION / PromethION platform from Oxford Nanopore in our laboratory; detailed culture conditions are described in the following text.

***S. cerevisiae* w303 culture:** *S. cerevisiae* strains w303 were cultured in Yeast Extract Peptone Dextrose (YPD) broth used as a complete medium for yeast growth. YPD medium, which contained 1 L of deionized water to 20 g bacto peptone, 10 g yeast extract, and 20 g dextrose, was sterilized by autoclaving for 20 min at 15 psi (1.05 $kg/cm^2$), and was stored at room temperature. Yeast cells were cultured at 30°C in a shaking incubator at 300 rpm for 24 to 36 hours.

***C. reinhardtii* culture:** High-quality genomic DNA was extracted from *C. reinhardtii* cultured under mixotrophic (constant light) or heterotrophic (constant dark) conditions in Tris-Acetate-Phosphate (TAP) medium during the pre-stationary phase. Samples of wild-type strain CC-1690 were placed in an intelligent temperature and illumination incubator under 4~6°C and 20~30 $\mu E/(m^2 \cdot s)$ light intensity. The naturally synchronized cells were induced using a 12 h/12 h light/dark cycle.

**Culture of O. sativa Japonica Group:** The seeds of O. sativa Japonica Group (Janponica Nipponbare) were sterilized, immersed in deionized water and germinated in the dark for 3 days. After germination, seedlings were transplanted into plastic pots filled with commercial substrate (PINDSTRUP, Denmark), and kept in a growth chamber at26/22° C ±1°C day/night temperature and light intensity of 600 $\mu molm^{-2}s^{-1}$. Four-weeks old seedlings were harvested for DNA isolation.

**Culture of retinoblastoma cell line WERI:** The human retinoblastoma cell line WERI was cultured in RPMI 1640 (Gibco Company, USA) supplemented with 20% fetal bovine serum (Biological Industries, USA). Cell cultures were incubated at 37°C and 5% $CO_2$, and media were replaced every 3~4 days. Cultures were maintained using centrifugation and resuspension in fresh medium, or media replacement after cell aggregates precipitated at the bottom of the flask. Cells were grown in suspension at a concentration of $10^5$~$10^6$ cells/ml.

### Supplementary Note 2: DNA extraction and purification

***S. cerevisiae* w303:** *S. cerevisiae* w303 cells were washed twice using phosphate-buffered saline (PBS) and collected by centrifugation at 4,000 rpm for 5 min. Samples were: (i) lysed in buffer with 1 ml lysozyme TLB and 20 µl RNase A (20 mg/ml), and then incubated for 1 h at 37°C; (ii) treated with 20 µl Proteinase K for 1.5 h at 50°C; (iii) purified with 1 volume phenol, 0.5 volume phenol-chloroform (1:1 by volume), 3 volume ice-cold absolute ethyl alcohol at 4,500 rpm for 10 min; (iv) washed in 80% ice-cold ethanol twice, collected by centrifugation (12,000 rpm, 15 min, 4°C), and eluted in 100 µl elution buffer(EB; 10 mMTris hydrochloride [pH 8.0]).

***C. reinhardtii* and *O. sativa Japonica Group*:** High-molecular-weight (HMW) DNA was isolated from *C. reinhardtii* cc1690 and *O. sativa Japonica Group* using the CTAB method. Briefly, about 0.2 g samples were re-suspended in 1 ml CTAB buffer containing 2% β-mercaptoethanol, incubated at 65°C for 30 min, and then centrifuged at 8,000 rpm for 5 min. The suspended nuclei were purified twice with chloroform-isoamyl alcohol (24:1 by volume) and once with 0.7 volume isopropyl alcohol at -20°C for 1 h. DNA precipitates were washed in ice-cold 75% ethanol twice, collected by centrifugation (12,000 rpm at 15 min and 4°C), dried under vacuum, and re-suspended in 100 ul EB[13] (10 mM Tris hydrochloride [pH 8.0]).

**Retinoblastoma cell line WERI:** 1 x $10^7$ frozen cells were lysed with 800 µl TEN Buffer, 100 µl 20% sodium dodecyl sulphate (SDS), and 100 µl proteinase K. This mixture was incubated at 56°C for 2 hours, purified with phenol-chloroform-isoamyl alcohol (25:24:1 by volume) and chloroform-isoamyl alcohol (24:1 by volume), and precipitated using 0.7 volume isopropyl alcohol at -20°C for 40 min. DNA precipitates were collected by centrifugation (12,000 rpm at 15 min and 4°C), washed twice in ice-cold 80% ethanol, dried under vacuum, re-suspended in 100 ul EB (10 mMTris hydrochloride [pH 8.0]), and combined with 2 µl RNase A (100 mg/ml) to cleave the RNA. To acquire high-quality DNA for the three datasets mentioned above, an additional purification step was performed using 0.8 volume magnetic beads from an AMPure XP kit (#A63882, Agencourt) according to the manufacturer’s instructions.

### Supplementary Note 3: Nanopore whole genome sequencing and base-calling

***S. cerevisiae* w303:** Sequencing libraries were constructed using a Ligation Sequencing Kit 1D (SQK-LSK108, Oxford Nanopore, UK) according to the manufacturer's instructions. Then, 5 µg high-molecular-weight genomic DNA was fragmented using g-TUBE (#520079, Covaris) centrifugation (conducted twice at 1,400 g for 2 min). Libraries were prepared according to the manufacturer's instructions. Briefly, NEBNext Ultra II End-Repair/dA-tailing module (#E7546, NEB) was used to end-repair and dA-tail the DNA fragments. Then, each dA-tailed sample was tethered to 1D adapter using NEBBlunt/TA Ligase Master Mix (#M0367, NEB). The prepared DNA library was loaded into R9.4 flow cells and sequenced on MinION sequencers (Oxford Nanopore). The raw data, collected in this experiment, were obtained as fast5 files after conversion of electrical signals into base calls via Albacore 1.1.0 (Oxford Nanopore Technologies).

***C. reinhardtii, O. sativa* and retinoblastoma cell line WERI:** Large insert-size libraries of *C. reinhardtii, O. sativa* and retinoblastoma WERI cells were created according to the manufacturer's protocols (Oxford Nanopore, UK). Briefly, 5 µg genomic DNA was sheared into ~20-30 kb fragments using g-TUBE (#520079, Covaris) centrifugation (twice at 1,400 g for 2 min) and size-selected (>8-10 kb) by Blue Pippin (Sage Science, MA) using a marker started at 5-12 min (0.75% DF Marker S1 High-Pass 6-10kb vs3) to ensure the removal of small DNA fragments. Genomic DNA libraries were prepared using a Ligation sequencing 1D kit (SQK-LSK109, Oxford Nanopore, UK). End-repair and dA-tailing of DNA fragments were performed using an Ultra II End Prep module (#E7546, NEB) according protocol recommendations. Each dA-tailed sample was tethered to 1D adapter using a Quick Ligation Module (#E6056, NEB). The prepared DNA library was loaded into a FLO-PRO002 flow cell and sequenced on PromethION sequencers (Oxford Nanopore, UK). The raw data collected in this experiment was obtained as fast5 files after conversion of electrical signals into base calls via guppy 2.0.8 (Oxford Nanopore, UK).

### Supplementary Note 4: Statistics for Nanopore datasets

To evaluate the performance of NECAT, we collected eight datasets for *E. coli, S. cerevisiae, A. thaliana*[14], *D. melanogaster*[15], *C. reinhardtii, O. sativa*, *S. pennellii*, and *H. sapiens*[16] (NA12878). Details can be found in Supplementary Table 1. Among these eight datasets, data on *E. coli, A. thaliana, D. melanogaster*, *S. pennellii*, and *H. sapiens* (NA12878) were available from public websites, and the other two datasets were generated using our in-house sequencing. Their corresponding short-reads datasets of Next Generation Sequencing (NGS) were collected from the related projects at NCBI. All SRA files were converted to fastq files using an SRA Toolkit[17] (https://www.ncbi.nlm.nih.gov/sra/docs/toolkitsoft/) from NCBI. Raw long-read files in fastq or fasta format were used as input files for these assembly pipelines. Nanopore fast5 format files and NGS fastq format files were used as input files for Nanopolish[4] and Pilon[5], respectively.

The results of basic statistical analysis for raw long reads (LRs) are shown in Supplementary Table 2. Seqkit (v0.8.0)[18] was used to directly calculate "Base Counts," "LR Count," "N50 Length," and "Mean Length." We then used scripts to calculate "N75 Length" and "N25 Length" based on results obtained using Seqkit. N75, N50, and N25 represented sequence lengths sorted in descending order when the accumulated length of the sequence reached 75, 50, and 25% of the total number of bases ("Base Counts"), respectively. Finally, we divided "Base Counts" by general genome size (*E. coli*: 4,600,000 base pairs [bp], *D. melanogaster*: 137,000,000 bp, *A. thaliana*: 125,000,000 bp, *S. cerevisiae*: 12,000,000 bp, *C. reinhardtii*: 120,000,000 bp, *O. sativa*:370,000,000, *S. pennellii:* 886,000,000 and *H. sapiens*: 3,000,000,000 bp) of the corresponding species to calculate coverage. Among these eight datasets, *A. thaliana* and *H. sapiens* datasets showed very low coverage (27X and 38X), while the other datasets showed more than 50X coverage.

N25 and N75 lengths were calculated using the following shell scripts:

```
ecoli=pathto/E.coli.fasta
yeast=pathto/w303.fastq
```

```
dro=pathto/dro.fastq
arab=pathto/arab.fastq
cre=pathto/cre.fastq
human=pathto/human.fastq
for i in ${ecoli} ${yeast} ${dro} $[19] ${cre} ${human};
do
seqkit fx2tab -j 10 -l -n -i -H ${i} | cut -f 4 | sed'1d' | sort -rn> ${i}.lenth.txt
seqkit stats -j 10 -a ${i} >>statistic.txt
all=$(awk'BEGIN{n=O}{n=n+$1}END{print n}' ${i}.lenth.txt)
echo"N75">>statistic.txt
awk'BEGIN{n=O}{if (n>="'$all'"*0.75){print $1;}n=n+$1;}' ${i}.lenth.txt | head
-n 1 >>statistic.txt
echo"N25">> statistic.txt
awk'BEGIN{n=O}{if (n>="'$all'"*0.25){print $1;}n=n+$1;}' ${i}.lenth.txt | head
-n 1 >> statistic.txt
done
```

**Supplementary Note 5：Error analysis of Nanopore raw reads**

Raw noisy LRs were corrected by mainstream consensus algorithms using the following steps: (1) building a multiple sequence alignment (MSA) from pairwise alignments and (2) choosing the correct base from MSA columns. FalconSense[20], Recon[6], Nanocorrect[4], and Dacoordare[21] are widely-used correction algorithms for Nanopore raw long reads. FalconSense uses the tagging and sorting approach to construct a consensus sequence based on consistent base-level and partial-order alignment. FalconSense and Recon were adopted for Nanopore sequence correction by fine-tuning parameters. Nanocorrect used a correction method similar to DAGCon[22], which encoded the MSA as a partial-order alignment with a directed acyclic graph (DAG). Dacoord resolves the corrected bases of repeated regions using a local de-Bruijn assembly-map algorithm. However, the accuracy and integrity of Nanopore corrected sequences produced by the above methods remained limited.

To determine whether the existing correction algorithms were feasible for correction of Nanopore raw reads, we needed to obtain the features of sequencing errors in Nanopore LR data. First, we analyzed error distribution of Nanopore datasets for *E. coli, S. cerevisiae*, *A. thaliana*, *D. melanogaster, C. reinhardtii*, *O. sativa*, *S. pennellii*, and *H. sapiens* (NA12878). We used reference genomes as standard sequences. Raw long reads of these Nanopore datasets were aligned using minimap2[23] against their corresponding reference genomes (Supplementary Table 1). Then, we statistically analyzed error distribution of each dataset according to the mismatched results.

Our results indicate that sequencing error rate of Nanopore reads was as high as 10-30% and broadly distributed（Figure 1A and Supplementary Table 3). We also found that the error rates of different positions differed broadly in each read, and the reads were generally present as high-error-rate subsequences (HERS), whose sequencing error rates were > 50% in these subsequences (Figure 1B). These sequencing error characteristics differed greatly from those in PacBio datasets (Figure 1). These results highlight the necessity of developing a specific consensus algorithm for Nanopore raw data.

We used the following scripts for aligning Nanopore datasets to their corresponding reference genomes:

```
minimap2 -t 20 -ax map-ont ${ref_fasta} ${reads_fasta} > ${species}_aln.sam
```

The error bases of all mapped reads were extracted and counted using the following scripts:

```
awk'{print $3"\t"$4"\t"$10"\t"$12}' ${species}_aln.sam
|awk'{split($4,a,":");print $1"\t"$2"\t"length($3)"\t"a[3]}' | awk'$3>100
{print $0}' | awk'/^NC/ {print $0}'> ${species}_stat_clean.txt
```

The distributions of sequencing errors for the six datasets were plotted using the following R scripts (Figure 1A):

```
ecolia=read.table(file="ecoli_stat_clean.txt")
yeast=read.table(file="yeast_stat_clean.txt")
arab=read.table(file="arab_stat_clean.txt")
dro=read.table(file="dro_stat_clean.txt")
yizao=read.table(file="yizao_stat_clean.txt")
human3=read.table(file="human_stat_clean.txt")
rice=read.table(file=" rice_stat_clean.txt")
tomato=read.table(file=" tom_stat_clean.txt")
for(i in 1:8)
{
if(i==1) {ecoli=ecolia} ; if(i==2) {ecoli=yeast} ; if(i==3) {ecoli=arab};
if(i==4) {ecoli=dro}; if(i==5) {ecoli=yizao}; if(i==6) {ecoli=human}; if(i==7)
{ecoli=rice}; if(i==8) {ecoli=tomato}
ecoli_n=numeric()
ecoli_s=cbind(ecoli,ecoli[,4]/ecoli[,3])
ecoli_t=dim(ecoli_s)[1]
for(j in 1:50){
  if(j==1){
    ecoli_n[1]=length(ecoli_s[ecoli_s[,5]<=0.01,5])/ecoli_t
  }
```

```
  else{
    pos2<-j/100
    pos1<-(j-1)/100
    ecoli_n[j]=length(ecoli_s[ecoli_s[,5]<=pos2 & ecoli_s[,5]>pos1,5])/ecoli_t
  }
}
if(i==1) {ecolin=ecoli_n}; if(i==2) {yeastn=ecoli_n};if(i==3) {arabn=ecoli_n}
if(i==4) {dron=ecoli_n}; if(i==5) {yizaon=ecoli_n}; if(i==6) {human3n=ecoli_n};
if(i==7) {ricen=ecoli_n}; if(i==8) {tomaton=ecoli_n}
}
pdf("read-error-distribution-fcraction.pdf")
plot(ecolin~c(1:50),ylab="Fraction  of  error  rate  (%)",xlab="Error  rate
(%)",ylim=c(0,0.20),col="darkgreen",type="l",axes=F,lwd=3,lty=1)
lines(yeastn~c(1:50),col="darkblue",lwd=3,lty=1)
lines(arabn~c(1:50),col="coral4",lwd=3,lty=1)
lines(dron~c(1:50),col="darkorange3",lwd=3,lty=1)
lines(yizaon~c(1:50),col="firebrick2",lwd=3,lty=1)
lines(human3n~c(1:50),col="yellow4",lwd=3,lty=1)
lines(ricen~c(1:50),col="chartreuse",lwd=3,lty=1)
lines(tomn~c(1:50),col="darkviolet",lwd=3,lty=1)
legend(25,0.18,legend=c("E.coli","Yeast","A.thaliana","D.melanogaster","C.re
inhardtii","Human"),col=c("darkgreen","darkblue" ,"coral4" ,"darkorange3","f
irebrick2","yellow4" ,"chartreuse","darkviolet"),lty=1,cex=1.5,box.lty=0,lwd=
2)
axis(2,at=c(0,0.05,0.10,0.15,0.20),labels=c("0","5","10","15","20"),las=1,lw
d=1,tick=T)
axis(1,at=c(0,10,20,30,40,50),labels=c("0","10","20","30","40","50"),las=1,l
wd=1,tick=T)
dev.off()
```

To further understand if there was a bias for sequencing errors among different genome positions, we calculated sequencing-error distribution for all mapped reads on different genome locations (Supplementary Figure 1). The scripts were:

```
awk'{print $1"\t"$2"\t"$2+$3"\t"$4/$3}' ${species}_stat_clean.txt |sort -k1,1
-k2,2n > ${species}.sorted.bed
refgenome=~/xsq/project/ONT_correct/distribution/data/${species}.fa
line=$(wc -l $refgenome |awk'{print $1}')
awk -v lin=$line '{if(NR!=lin&&/^>/) {print $1"\t"NR;tmp=$1}
if(NR==lin) {print tmp"\t"NR}}' $refgenome\
|awk'NR==1 {tmp1=$1;tmp2=$2 }
NR!=1 {print tmp1"\t"tmp2"\t"$2"\t"($2-tmp2-1)*80; tmp1=$1; tmp2=$2}'
|awk'{len=$4/10000; for(i=1;i<=len;i++) {print
$1"\t"(i-1)*10000+1"\t"10000*i}}' |awk'{split($1,a,">"); print
a[2]"\t"$2"\t"$3 }'>${species}_10000.bed
export PATH=$PATH:/software/bedtools2/bin
bedtools intersect -loj -a ${species}_10000.bed -b
${species}.sorted.bed>position_{species}.txt
```

To further understand the different error distribution in each read, we extracted each raw read, and calculated the mismatch and indel base number in a region having a length >500 bp (Figure 1B). The scripts were:

```
awk '{tmp=0; for (i=1;i<length($6);i++) {st=substr($6,i,1); if(st~/[0-9]/)
{ss=ss""st}
   if(st=="M") {mn=mn+ss;mi=mi+ss;tn=tn+ss;ss=""}
   if(st=="D") {tn=tn+ss;ss=""}
   if(st=="I") {tn=tn+ss;mi=mi+ss;ss=""}
   if(st~/[A-Za-z]/ ) {ss=""}
   if(mi>500&&st=="M")
{for(j=mi;j>500;j=j-500){re=re"_"(mn+500-j)/tn;tn=j-500;mn=j-500;mi=j-500}}
if(mi>500&&st=="I")
```

```
{for(j=mi;j>500;j=j-500){re=re"_"mn/tn;tn=j-500;mi=j-500;mn=0 }}
if(mi==500){re=re"_"mn/tn;tn=0;mi=0;mn=0}
}; print re}'  ERR2173373.21178.sam | sed s/_/'\n'/g | awk 'NR>1{print
(NR-2)*500"-"(NR-1)*500"\t"1-$1}' > stat_500.res
```

Then, error subsequences of 500 bp in each read were plotted and beautified by Excel and Adobe Illustrator.

The high error rate subsequences (HERS) of eight datasets were extracted as similarly as each read. The scripts were:

```
awk  -v  var=10  'length($10)>var*1000  {tmp=0;  for  (i=1;i<length($6);i++)
{st=substr($6,i,1); if(st~/[0-9]/) {ss=ss""st}
     if(st=="M") {mn=mn+ss;tn=tn+ss;ss=""}
      if(st=="D") {tn=tn+ss;ss=""}
     if(st=="I") {tn=tn+ss;ss=""}
     if(st~/[[:alpha:]]/ ) {ss=""}
     if(tn>500) {if(mn/tn<0.50) {tmp=1;break}; mn=0;tn=0 }
    }if(tmp==1) {tsum=tsum+1;;mn=0;tn=0};
     print NR,length($10),tmp}
    '  mutilsam/tom_raw_aln$i.sam > stat_500.res.txt
```

After extracting high error rate subsequences, HERS distributions for the six datasets were plotted using the following R scripts (Figure 1C):

```
ecoli=read.table(file="ecoli/stat_500.res.txt")
yeast=read.table(file="yeast/stat_500.res.txt")
arab=read.table(file="arab/stat_500.res.txt")
dro=read.table(file="dro/stat_500.res.txt")
yizao=read.table(file="yizao/stat_500.res.txt")
human=read.table(file="human/stat_500.res.txt")
rice=read.table(file=”rice/stat_500.res.txt”)
tomato=read.table(file=tomato/stat_500.res.txt”)
result=matrix(,8,41)
for(j in 1:8)
```

```
{if(j==1)  {a=ecoli}  ; if(j==2){a=yeast}; if(j==3){a=arab}; if(j==4){a=dro};
if(j==5){a=yizao}; if(j==6){a=human}; if(j==7){a=rice}; if(j==8){a=tomato};
 for(i in 1:41)
 {  usum=a[a[,2]>=(i+9)*1000&a[,3]==1,3]
    tsum=a[a[,2]>=(i+9)*1000,3]
    if (length(usum)>=500){ result[j,i]=length(usum)/length(tsum)}}
}
tmp1=result[1,][!is.na(result[1,1:41])];tmp2=result[2,][!is.na(result[2,1:41
])];tmp3=result[3,][!is.na(result[3,1:41])];tmp4=result[4,][!is.na(result[4,
1:41])];tmp5=result[5,][!is.na(result[5,1:41])];tmp6=result[6,][!is.na(resul
t[6,1:41])];tmp7=result[7,][!is.na(result[7,1:41])];tmp8=result[8,][!is.na(r
esult[8,1:41])]
pdf("HER length.pdf")
plot(result[1,1:length(tmp1)]~c(1:length(tmp1)),type="l",axes=F,lwd=3,lty=1,
col="darkgreen", ylim=c(0,0.5), xlim=c(1,41), xlab="Read length(kb)", ylab=
"Fraction of reads with HER")
axis(2,at=c(0,0.1,0.20,0.3,0.4,0.5),labels=c("0","10","20","30","40","50"),
las=1,lwd=1,tick=T)
axis(1,at=c(1,11,21,31,41),labels=c("10","20","30","40","50"),las=1,lwd=1,ti
ck=T)
lines(result[2,1:length(tmp2)]~c(1:length(tmp2)),col="darkblue",lwd=3,lty=1)
lines(result[3,1:length(tmp3)]~c(1:length(tmp3)),col="coral4",lwd=3,lty=1)
lines(result[4,1:length(tmp4)]~c(1:length(tmp4)),col="darkorange3",lwd=3,lty
=1)
lines(result[5,1:length(tmp5)]~c(1:length(tmp5)),col="firebrick2",lwd=3,lty=
1)
lines(result[6,1:length(tmp6)]~c(1:length(tmp6)),col="yellow4",lwd=3,lty=1)
lines(result[7,1:length(tmp7)]~c(1:length(tmp7)),col="chartreuse",lwd=3,lty=
1)
lines(result[8,1:length(tmp8)]~c(1:length(tmp8)),col="darkviolet",lwd=3,lty=
```

```
1)
legend(3,0.5,legend=c("E.coli","Yeast","A.thaliana","D.melanogaster",
"C.reinhardtii","Human"),col=c("darkgreen","darkblue" ,"coral4" ,"darkorange
3","firebrick2", "yellow4" ,"chartreuse","darkviolet"),lty=1,cex=1.2,box.lty
=0,lwd=2)
dev.off()
```

### Supplementary Note 6: Performance of error correcting algorithms

Due to the high sequencing error discrepancy between Nanopore raw reads and PacBio raw reads (Figure1 and Supplementary Note 5), the existing correction methods developed specifically for PacBio reads are unsuitable for Nanopore data. To date, there is no correction method that fully accounts for characteristics of sequencing errors occurring in Nanopore data.

In this study, we developed a novel progressive two-step error correction algorithm called NECAT with adaptive candidate-read selection for Nanopore raw reads. In order to validate the rationality and reliability of our novel algorithm, we examined the performance of NECAT in correcting the eight datasets described above (Supplementary Table 1). For comparison, we also evaluated the accuracy of reads corrected by Canu[24], another widely-used correction tool for Nanopore raw reads. Specifically, for each dataset, we calculated error rates of: the raw dataset, corrected reads after step one in NECAT, corrected reads after step two in NECAT, and reads corrected by Canu[24]. For this, we mapped the four datasets to the reference using minimap2[23] as described in Supplementary Note 5. Then, results of the alignment were used to calculate error distribution. Error rates were grouped by 1, 2, 3, 4, 5, 6, 7, 8, 9, 10, 10-15, 15-20, 20-25, 25-30, and 30-100%, and results are listed in Supplementary Table 4. The following scripts were used:

```
for i in 1 2 "canu" "raw"
do
cd $i
awk'{print $3"\t"$4"\t"$10"\t"$12}' ${species}_${i}_aln.sam |awk'{split($4, a,
":"); print $1"\t"$2"\t"length($3)"\t"a[3]}' | awk'$3>100 {print $0}' |
awk'/^chr/ {print $0}'> ${species}_stat_clean_$i.txt
Rscript correct_stats_ref.r ${species}_stat_clean_$i.txt "correct_stat.result"
cd ..
done
```

In each raw dataset, we then analyzed a HERs region having a length >500 bp. For mapped reads in each of the four datasets, we evaluated raw reads, corrected reads after first correction of NECAT, corrected reads after second correction of NECAT, and corrected read output by Canu. Considering canu only selects the longest 40x for correction by default, we extracted the sub-dataset with equal coverage from the raw dataset, corrected reads after step one in NECAT and corrected reads after step two in NECAT. The scripts were:

```
###species can use eight species, we take e.coli for example
species=ecoli
size=`ls -ltr ecoli_canu.fasta | awk '{print $5}'`
for i in 1 2 "raw"seqfasta= ecoli_$i.fasta
awk 'NR%2==1 {tmp=$1}NR%2==0 {print tmp"_XSQ_"$0"\t"length($0)}'\ ${seqfasra}
| sort -nr -k 2 | awk -v si=$size 'tmp=tmp+$2\
 {if(tmp<si){print $0} if(tmp>=si) exit}' | \
awk '{split($1,a,"_XSQ_");print a[1]"_XSQ_"$2"\n\r"a[2]}' >
rice${i}_filter.fasta
```

In order to calculate the number of gaps, we generated alignment paf files using minimap2. The scripts were:

```
reffasta=ecoli_k12_genomic.fna
for i in 1 2 "raw"
do
echo $i
mkdir -p ~/alignment/minimap2/$species/$i
cd~/alignment/minimap2/$species/$i
seqfasta= /data/$i/ecoli$i_filter.fasta
minimap2 -t 20 -x map-ont ${reffasta} ${seqfasta} >${species}_${i}_aln.paf
done
minimap2 -t 20 -x map-ont ${reffasta} ecoli_canu.fasta > ecoli_canu_aln.paf
```

For raw reads, we extracted all the reads with gaps >500 bp, and counted the number of HERs regions using the following scripts:

```
awk'{print $6"_"$1"\t"$3"\t"$4"\t"$2}' ${species}_raw_aln.paf> \
${species}_raw_bed.txt
sort -k1,1 -k2,2n ${species}_raw_bed.txt |uniq>in.sorted.bed
bedtools merge -iin.sorted.bed -d 500 | awk'{print $1}' |uniq -d
|awk'{split($1,a,"_"); {print a[3]"\t"1"\t"10000}}'>
${species}_gap_read_name.txt
wc -l  ${species}_gap_read_name.txt
```

For these raw reads with gaps, we re-calculated the HERs region number in these reads after first correction of NECAT, after second correction of NECAT, and after correction of Canu. For outputted corrected reads from Canu, we extracted the reads having a HERs region >500 bp and counted the number of these regions using the following scripts:

```
awk'split($1,a,"_") {print a[1]"\t"$3"\t"$4"\t"$6}' ${species}_canu_aln.paf>
${species}_canu_bed.paf
sort -k1,1 -k2,2n ${species}_gap_read_name.txt |uniq |bedtools merge -i - -d 500
|awk'{print $1}' |uniq -d >read_gap.result.final
wc -l read_gap.result.final
```

For corrected reads produced by step one and step two in NECAT, reads having a HERs region >500 bp were extracted using the following scripts:

```
for i in "ecoli" "yeast" "dro" "ara" "yizao" “human” “rice” “tomato”
do
for j in 1 2
do
cd ${i}/${j}
awk'{split($1,a,"_\\(");print a[1]"\t"$3"\t"$4"\t"$6}' ${i}_${j}_aln.paf>
${i}_${j}_bed.paf
cd ../..
done
done
```

Finally, the gap number was counted by:

```
sort -k1,1 -k2,2n ${species}_1_bed.paf |uniq |bedtools merge -i - -d 500
|awk'{print $1}' |uniq -d |wc -l
```

The number of HERS regions with large gaps > 500 bp in each raw and corrected dataset can be found in Table 1.

**Supplementary Note 7: Comparison of assembly pipelines**

We compared the quality of assembly results and running time for Canu (v1.8)[24], Canu (v1.8)+smartdenovo (5cc1356b)[25], Smartdenovo (5cc1356b), miniasm (1552e6f9), wtdbg2 (v2.5), Flye (2.6), and NECAT pipelines. Running time was recorded from the log files. All assemblers ran on a 4-core 24-thread Intel(R) Xeon(R) 2.4 GHz CPU (CPU E7-8894[v4]) machine with 3 TB of RAM; the OS was Centos 7.3 64-bit (Linux). The eight datasets (*E. coli, S. cerevisiae, A. thaliana*, *D. melanogaster*, *C. reinhardtii*, *O. sativa*, *S. pennellii* and *H. sapiens*) composed of Nanopore long reads were assembled by the pipelines. The *de-novo* genome assemblies of eight datasets and results of statistical analyses are shown in Table 2 and Supplementary Table 7.

Canu pipeline was run as:

```
echo Start: $(date "+%Y-%m-%d %H:%M:%S")
canu -p $genomeName -d $genomeName genomeSize=$genomeSize maxMemory=1000
maxThreads=$threads useGrid=false -nanopore-raw input.fastq
echo End: $(date "+%Y-%m-%d %H:%M:%S")
```

where *$genomeName* was set to *E. coli, S. cerevisiae, A. thaliana*, *D. melanogaster*, *C. reinhardtii*, *O. sativa* and *S. pennellii,* respectively, and *$genomeSize* was set to 4.8M, 13M, 130M, 130M, 120M, 400 M and 1G, respectively. $theads was set to 32 for *E. coli, S. cerevisiae, A. thaliana*, *D. melanogaster* and *C. reinhardtii* and 64 for *O. sativa* and*S. pennellii.*

For Canu+smartdenovo pipeline, the output file *$genomeName.correctedReads.fasta* from the Canu pipeline was used as input file to the Canu+smartdenovo pipeline; the script was as follows:

```
echo Start: $(date "+%Y-%m-%d %H:%M:%S")
smartdenovo.pl –p $genomeName -t $threads -c 1 $genomeName.correctedReads.fasta >
$genomeName.mak
make -f $genomeName.mak
```

```
echo End: $(date "+%Y-%m-%d %H:%M:%S")
```

For the Flye pipeline, we used the following script:

```
echo Start: $(date "+%Y-%m-%d %H:%M:%S")
flye --nano-raw input.fastq --out-dir $genomeName --genome-size $genomeSize
--threads $threads
echo End: $(date "+%Y-%m-%d %H:%M:%S")
```

Flye failed to run on raw reads of *E. coli* and *C. reinhardtii*, for the input files contained malformated reads and duplicate reads. We used the following scripts to filter the raw reads before running Flye. For *E. coli*, the script was:

```
fsa_rd_tools longest --base_size 0 --discard_illegal_read --ifname inputfile
--ofname outputfile
```

`fsa_rd_tools` was a tool in NECAT pipeline.

For *C. reinhardtii*, the script was:

```
python3 remove_dup_name.py inputfile outputfile
```

`remove_dup_name.py` contained following code:

```
import sys
from collections import defaultdict
from Bio import SeqIO
ifname = sys.argv[1]   # xxx.fasta or  xxx.fastq
ofname = sys.argv[2]
names = defaultdict(int)
with open(ofname, "w") as ofile:
    for i, rec in enumerate(SeqIO.parse(ifname, ifname[-5:])):
        names[rec.id] += 1
        if names[rec.id] == 1:
            SeqIO.write(rec, ofile, ofname[-5:])
```

wtdbg2 pipeline was ran as

```
echo Start: $(date "+%Y-%m-%d %H:%M:%S")
wtdbg2.pl -t $threads -x ont -g $genomeSize -o $genomeName input.fastq
echo End: $(date "+%Y-%m-%d %H:%M:%S")
```

Smartdenovo pipeline was ran as:

```
awk 'NR%4==1||NR%4==2' all.fastq | sed 's/^@/>/g' > reads.fa
echo Start: $(date "+%Y-%m-%d %H:%M:%S")
smartdenovo.pl -p $genomeName -t 32 -c 1 reads.fa > dro_smart.mak
make -f dro_smart.mak
echo End: $(date "+%Y-%m-%d %H:%M:%S")
```

miniasm pipeline was ran as:

```
echo Start: $(date "+%Y-%m-%d %H:%M:%S")
minimap2 -x ava-ont -t32 all.fastq all.fastq | gzip -1 > reads.paf.gz
miniasm -f all.fastq reads.paf.gz > $genomeName.gfa
awk '/^S/{print ">"$2"\n"$3}' $genomeName.gfa | seqkit seq > $genomeName.fasta
echo End: $(date "+%Y-%m-%d %H:%M:%S")
```

NECAT pipeline first generated configuration file (necat_cfg.txt), as shown below:

```
PROJECT=$genomeName
THREADS=$threads
ONT_READ_LIST=read_list.txt
GENOME_SIZE=$genomeSize
MIN_READ_LENGTH=3000
PREP_OUTPUT_COVERAGE=40
OVLP_FAST_OPTIONS="-n 500 -z 20 -b 2000 -e 0.5 -j 0 -u 1 -a 1000"
OVLP_SENSITIVE_OPTIONS="-n 500 -z 10 -e 0.5 -j 0 -u 1 -a 1000"
CNS_FAST_OPTIONS="-a 2000 -x 4 -y 12 -l 1000 -e 0.5 -p 0.8 -u 0"
CNS_SENSITIVE_OPTIONS="-a 2000 -x 4 -y 12 -l 1000 -e 0.5 -p 0.8 -u 0"
TRIM_OVLP_OPTIONS="-n 100 -z 10 -b 2000 -e 0.5 -j 1 -u 1 -a 400"
```

```
ASM_OVLP_OPTIONS="-n 100 -z 10 -b 2000 -e 0.5 -j 1 -u 0 -a 400"
NUM_ITER=2
CLEANUP=1
USE_GRID=false
GRID_NODE=0
SMALL_MEMORY=0
CNS_OUTPUT_COVERAGE=30
FSA_OL_FILTER_OPTIONS=""
FSA_ASSEMBLE_OPTIONS=""
FSA_CTG_BRIDGE_OPTIONS=""
```

Then, it was run as:

```
echo Start: $(date "+%Y-%m-%d %H:%M:%S")
necat.pl bridge necat_cfg.txt
echo End: $(date "+%Y-%m-%d %H:%M:%S")
```

The read_list.txt contained the path of corresponding sequencing data; *$genomeName* was set to *E. coli, S. cerevisiae, A. thaliana*, *C. reinhardtii*, *D. melanogaster*, *O. sativa, S. pennellii* and *H. sapiens*, respectively, and *$genomeSize* was set to 4,800,000, 13,000,000, 130,000,000, 130,000,000, 120,000,000 400,000,000, 1,000,000,000 and 3,000,000,000, respectively.

For large genomes, NECAT used more corrected reads to obtain more robust assemblies. Therefore, we adjusted the parameters for *O. sativa* as shown below:

```
CNS_OUTPUT_COVERAGE=40
FSA_OL_FILTER_OPTIONS="—min_coverage 3"
```

And we adjusted the parameters for *S. pennellii* as shown below:

```
CNS_OUTPUT_COVERAGE=40
```

We also adjusted the parameters for *H. sapiens* and WERI as shown below:

```
MIN_READ_LENGTH=500
PREP_OUTPUT_COVERAGE=
```

```
OVLP_FAST_OPTIONS="-n 200 -z 10 -b 2000 -e 0.5 -j 0 -u 1 -a 400"
OVLP_SENSITIVE_OPTIONS="-n 200 -z 10 -e 0.5 -j 0 -u 1 -a 400"
CNS_FAST_OPTIONS="-a 400 -x 4 -y 12 -l 500 -e 0.5 -p 0.8 -u 0"
CNS_SENSITIVE_OPTIONS="-a 400 -x 4 -y 12 -l 500 -e 0.5 -p 0.8 -u 0"
CNS_OUTPUT_COVERAGE=45
```

### Supplementary Note 8: Validation of the WERI genome

The new WERI assembly from Nanopore data was polished four times using the same scripts as those shown in **Supplementary Note 13**. We compared the WERI assembly against human reference genome hg38. The newly-assembled genome was aligned to the reference genome, and the Mummer plot between them was generated using MUMmer (v4.0)[7] with the following script (**Supplementary Figure 2**):

```
nucmer --mum -l 10 -c 1000 --banded ${ref.fasta} ~/project/weri/ONT_asm.fasta
dnadiff -d out.delta
mummerplot out.delta --fat -f -png
```

Because MUMmer was operated using a unique anchor matching option to accelerate the alignment, some repetitive sequences remained unaligned. The entire process of alignment and figure generation can be reproduced using scripts available on the MHAP home page[20] (assuming that Perl, Python, and MUMmer[7] are placed in the correct path), and by running the script below; this generates a figure designated as asm.pdf (**Figure 3**).

```
sh makeHuman.sh ref.fasta asm.fasta
```

Based on out.rdiff file output by dnadiff, structural differences (>10 bp) were extracted using the following scripts:

```
awk ' {if($3<=$4&&$7*$7>100) print $1 "\t" $3"\t"$4"\t"$2"\t"$7
if($3>$4&&$7*$7>100) print $1 "\t" $4"\t"$3"\t"$2"\t"$7 }' ./out.rdiff >
weri_10.bed
wc -l weri_10.bed
```

We then used a custom script to convert the SV regions in the WERI assembly genome to the reference hg38:

```
awk '{if($3>$4&&$7*$7>100) print
$1"\t"$4"\t"$3"\t"$2"\t"$7"\t"($3-1)"\t"($4+1)
if ($3<=$4&&$7*$7>100) print
```

```
$1"\t"$3"\t"$4"\t"$2"\t"$7"\t"($3-1)"\t"($4+1)}' ./out.qdiff > hg38_gap.tsv
awk '{if($3>$4&&$7*$7>100) print
$1"\t"$4"\t"$3"\t"$2"\t"$7"\t"($3-1)"\t"($4+1)
if ($3<=$4&&$7*$7>100) print
$1"\t"$3"\t"$4"\t"$2"\t"$7"\t"($3-1)"\t"($4+1)}'  ./out.rdiff > weri_gap.tsv
python3 query.py  –c hg38_gap.tsv –w weri_gap.tsv –a out.1coords > weri2hg38.tsv
```

To validate SV regions detected in WERI, we re-aligned the original sequencing data with SV regions±1000 bp. SV regions were extracted with:

```
awk '{if($5>=$6) print $4,":",$6-1000,"-",$5+1000
if($5<$6) print $4,":",$5-1000,"-",$6+1000}' weri2hg38.tsv | sed 's/ //g' > qgap
for i in $(cat qgap);do samtools faidx ./ref.fasta $i >> all_gap.fasta;done
# re-align the raw nanopore reads to all_gap.fasta
minimap2 -x map-ont -t $NPROC ./all_gap.fasta ./fq > all_gap.paf
```

Then, we calculated the number of SV regions with read coverage:

```
awk '($8<=($7-1000))&&($9>=1000){print $6,"\t",$7,"\t",$8,"\t",$9}'
all_gap.paf > real_map
awk '{print $1}' real_map | sort | uniq -c | tee real_map_list | wc l
awk '{split($2,a,"\[:-\]");print
a[1],"\t",(a[2]+1000),(a[3]-1000),$1}' ./real_map_list > real_map_list_raw
#generate merge.tsv
cat ./real_map_list_raw | xargs -n 4 -P 10 ./merge.sh
# generate merge.bed
awk '{if($2>$3){print $1,$3,$2,$4}else{print $1,$2,$3,$4}}' ./merge.tsv | sed
's/ /\t/g'  | grep -v 'chrY' > merge.bed
```

We also aligned the raw nanopore long reads and Illumina short reads to human reference genome hg38, and used Sniffles[26] and Lumpy_sv[27] to call SVs in mapping results using the scripts shown below:

*Sniffles:*

```
export PATH=/ /software/Sniffles-1.0.10/bin/sniffles-core-1.0.10:$PATH
export PATH=/ /software/ngmlr-0.2.7:$PATH
ngmlr -t $NPROC -r $refsequence -q $fq -o reads.sam -x ont
samtools view -bS reads.sam | samtools sort -@ $NPROC - -o reads.sorted.bam
sniffles -t $NPROC -m reads.sorted.bam -v tgs.weri.vcf

Lumpy_sv:
bwa  mem  -R  "@RG\tID:id\tSM:sample\tLB:lib"  reference.fasta  sample.1.fq
sample.2.fq  |  samblaster  --excludeDups  --addMateTags  --maxSplitCount  2
--minNonOverlap 20 | samtools view -S -b -  >  sample.bam
samtools view -b -F 1294 sample.bam
| samtools sort  -o sample.discordants.sorted.bam
samtools view -h sample.bam \
  | scripts/extractSplitReads_BwaMem -i stdin \
  | samtools view -Sb - \
  | samtools sort  -o sample.splitters.sorted.bam
lumpyexpress \
    -B sample.bam \
    -S sample.splitters.bam \
    -D sample.discordants.bam \
    -o output.vcf
export PATH= /software/VCFtools/bin:$PATH
cat ngs.weri.vcf | vcf-sort > sorted.ngs.vcf
cat tgs.weri.vcf | vcf-sort > sorted.tgs.vcf
bzip sorted.ngs.vcf
bzip sorted.tgs.vcf
bcftools stats ./sorted.ngs.vcf.gz > ngs.stat
bcftools stats ./sorted.tgs.vcf.gz > tgs.stat
#index
tabix -p vcf sorted.ngs.vcf.gz
```

```
tabix -p vcf sorted.tgs.vcf.gz
# generate 0000.vcf  0001.vcf  0002.vcf  0003.vcf
-l
# weri SV and ngs
bedtools intersect -a ./merge.bed -b ./sorted.ngs.bed -wa -loj | awk
'$5!="."{print}' | wc -l
# weri SV, ngs and tgs overlap
bedtools intersect -a ./merge.bed -b ./comm.ngs2tgs.bed -wa | wc -l
bcftools isec sorted.ngs.vcf.gz sorted.tgs.vcf.gz -p ./
# convert vcf to bed
awk '{split($8,a,"RE=");print $1,$2,($2+1),a[2]}' ./sorted.tgs.vcf | grep -v '#'
| grep -v 'chrY' | sed 's/ /\t/g'> sorted.tgs.bed

awk '{split($10,a,":");print $1,$2,($2+1),a[2]}' ./sorted.ngs.vcf | grep -v '#'
| grep -v 'chrY' | sed 's/ /\t/g' > sorted.ngs.bed
echo "CHROM POS ID REF ALT QUAL FILTER Coverage" > 0002.head
awk '{split($10,a,":");print $1,$2,$3,$4,$5,$6,$7,a[2]}' ./0002.vcf | grep -v
'#' | cat 0002.head - > ngs.commom.add_cov.vcf
grep -v CHROM ./ngs.commom.add_cov.vcf | awk '{print $1,$2,($2+1)}' | sed 's/
/\t/g' > comm.ngs2tgs.bed
# weri SV and tgs
bedtools intersect -a ./merge.bed -b ./sorted.tgs.bed -wa -loj | awk
'$5!="."{print}' | wc -l
```

### Supplementary Note 9: Overlap-filtering strategy

Overlap-filtering is critical in genome assembly. High-error-rate overlaps introduce errors and complicate assembly. Conversely, an overly strict filtering strategy can reduce contiguity of the results. Error distribution of sequencing data varies greatly. In order to adapt to different data, we adopted a heuristic filtering strategy to remove high-error-rate overlaps. Two metrics, the identity obtained by dividing length of the overlap by the number of matching bases, and the overhang that is the distance of an overlap from the 5' or 3' end of the read, are used to identify high error rate overlaps.

First, we examined overlap identities. For each read, we collect its overlaps and compute the mean of identities of the overlaps as its identity. After obtaining all read identities, we computed the weighted median ($m_g^{id}$) and weighted median absolute deviation ($MAD_g^{id}$) of them, where the weight is the read length. We used the following formula to calculate global threshold of overlap identity ($th_g^{id}$):

$$th_g^{id} = \min(m, m_g^{id}) - n * k * MAD_g^{id}. \#(1)$$

Here $k$ is equal to 1.4862, a constant scale factor multiplied by *MAD* to obtain an estimation of the standard deviation $\sigma$. According to our experience, $m$ and are $n$ are set to 0.98 and 6, respectively. After obtaining global threshold for overlap identity, we calculated the local threshold. For each read, we accumulate the lengths of its overlaps. If the sum was less than $\max(c_{min}, 0.5 * c) * l$, where $c_{min}$ is a user-set parameter (default value is 25), $c$ is the coverage of corrected reads, and $l$ is read length, we set the local threshold $th_l^{id}$ to global threshold $th_g^{id}$, because the data were too small to show statistical significance. Otherwise, we sorted the overlaps in descending order according to the product of overlap identity and overlap length. We collected the first several overlaps in which the sum of their lengths was no more than $\max(2 * c_{min}, 1.5 * c) * l$. Then, we computed weighted median ($m_l^{id}$) and weighted median absolute deviation ($MAD_l^{id}$) of these overlaps, where the weight was overlap length. Local identity threshold $th_l^{id}$is was set to $\max\left(th_g^{id}, \left(\min(m, m_l^{id}) - n * k * MAD_l^{id}\right)\right)$, where *m* and *n* were set to 0.99 and 6 by default. Next, we used $th_l^{id}$ to filter out the read overlaps. If overlap identity was less than $th_l^{id}$, the overlap was removed.

We used a similar process to assess read overhang in the overlaps. For each read, we collected the maximum of its overhangs. Then, we computed the weighted median ($m_g^{oh}$) and weighted median absolute deviation ($MAD_g^{oh}$), where weight was read length. The formula used to calculate global threshold of an overhang is provided in (2), where *m* and *n* were set to 30 and 6 by default, respectively.

$$th_g^{oh} = \max\left(m, m_g^{oh}\right) + n * k * MAD_g^{oh}. \#(2)$$

Next, we collected read overhangs at the 5’ or 3’ end separately, and computed the local thresholds for them. For each end, if the number of read overhangs is less than $max(c_{min}, 0.5 * c)$, then $c_{min}$ is a user-set parameter (default value is 25), $c$ is coverage of corrected reads, and local threshold $th_l^{oh}$ is set to global threshold $th_g^{oh}$. Otherwise we sorted overhangs in ascending order according to results obtained by dividing the overlap overhang by overlap length. We collected the first several overhangs, the number of whom is no more than $\max(2 * c_{min}, 1.5 * c)$. Then, we computed weighted median ($m_l^{oh}$) and weighted median absolute deviation ($MAD_l^{oh}$)

for these overhangs, where weight was overlap length. Local identity threshold $th_l^{oh}$ was set to $\min\left(th_g^{oh}, \left(\max\left(m, m_l^{oh}\right) + n * k * MAD_l^{oh}\right)\right)$, where $m$ and $n$ were set to 10 and 6 by default, respectively. The overlap was removed if the read overhang at 5' or 3' end was greater than $th_l^{oh}$ of the corresponding end.

In addition to assessing overlap identity and read overhang, we used the following filtering strategies.

1. We calculated the coverage for each base in the reads according to overlaps between them. For each read, we obtained three metrics, minimum coverage of all bases ($c_{min}$), maximum coverage of all bases ($c_{max}$), and the difference between minimum coverage and maximum coverage ($c_{diff}$). The procedure used three thresholds, designated as *min_coverage, max_coverage*, and *max_diff_coverage*, to assess read metrics and automatically select thresholds based on statistical results. If $c_{min}$ is less than *min_coverage*, $c_{max}$ is larger than *max_coverage*, and $c_{diff}$ is larger than *max_diff_*coverage, the reads and related overlaps are removed. Our analysis of the yeast dataset indicated that *min_coverage* should be set to the first value not exceeding 30% of the value for the first trough of the histogram of all $c_{min}$s, as shown in Supplementary Figure 15. We suggest that *max_coverage* and *max_diff_coverage* be set to (100-*x*)-th percentile (*x*=0.01 by default). These thresholds can also be specified by users. After some of the reads are filtered out, coverages of each read may change. This filtering strategy is executed twice to increase robustness of the results.

2. In this step, we assessed the overlaps and counted the number of reads having an overlap with the first read and covering the 5'- or 3'-end of the second read. If the number was less than *min_coverage* - 1, this overlap was filtered out.

3. The contained reads and related overlaps were filtered out.

4. Finally, for each read, we sorted the overlaps covering its 5'- or 3'-end by aligned length, respectively. The best overlaps can be selected using *bestn*, a parameter specified by users.

### Supplementary Note 10: Genome polishing and assembly validation

Different polishing strategies were used for different genome-assembly pipelines (NECAT, Canu[24], and Canu+smartdenovo[25]) and different species (*E. coli, S. cerevisiae, A. thaliana*, *D. melanogaster*, *C. reinhardtii*, *O. sativa* and *S. pennellii*):

1. Nanopolish (v0.10.2)[4] was used to further polish the genome using fast5 files and corresponding fasta/fastq files. Finally, the genome was polished three times using NGS data with Pilon (v1.22)[5] and generated the final genome.

2. For the *A. thaliana*, we used the Arrow in smrtlink (v5.1.0)[28] to polish the draft genome with Sequel Bam files because the raw fast5 files required by Nanopolish were not available.

3. For the *O. sativa* and Human, we used minimap2[23] (v2.10-r761) with "-x map-ont" and Racon[6] (v1.3.1) with default parameters to polish the draft genome four times using raw reads.

Alignments and validation results of statistical analysis are shown in Supplementary Figures 6-13.

We then mapped the assembled genomes onto their reference genomes, and counted single-nucleotide polymorphisms (SNPs) and large indels using dnadiff[29] and GEGE[30]. The five genome assemblies for *E. coli*, *S. cerevisiae*, *A. thaliana*, *D. melanogaster* and *C. reinhardtii* and were aligned to their reference genomes and plotted using MUMmer (v4.0)[7]. Results were generated using the following scripts:

```
nucmer --mumreference -l 100 -c 1000 -d 10 --banded -D 5 ${ref.fasta} ${asm.fasta}
delta-filter -i 95 -o 95 out.delta> out.best.delta
dnadiff -d out.best.delta
mummerplotout.best.delta --fat -f -png
```

We also compared genome assemblies for *E. coli, S. cerevisiae, A. thaliana, C. reinhardtii*, and *D. melanogaster* generated using Canu, Canu+Smartdenovo, Smartdenovo, miniasm, wtdbg2, Flye, and NECAT pipelines. The following scripts were used to evaluate Indel gaps in these genome assemblies:

```
awk'{if($2=="GAP"&&sqrt($7*$7)>=10) {print $0 }}' ${out.qdiff} > indelM10.txt
awk'{if($2=="GAP"&&sqrt($7*$7)<10) {print $0 }}' ${out.qdiff} > indelL10.txt
```

SNPs and indels between the assembly genome and reference genome are listed in Supplementary Table 8.

### Supplementary Note 11: Analysis of repeat regions in *D. melanogaster*

Repeat regions are one of the greatest challenges in genome assembly. To assess transposable_element (TE)[12] resolution in NECAT assembly, we analyzed the TE repeat families and aligned the annotated *D. melanogaster* genome[11] to the seven assembled contigs from the genome assemblies pipelines (Canu, canu+smartdenovo, Smartdenovo, miniasm, wtdbg2, Flye, and NECAT). Genome FlyBase 5.57_FB2014_0310 was downloaded from:

[ftp://ftp.flybase.net/genomes/dmel/dmel_r5.57_FB2014_03/fasta/dmel-all-gene-r5.57.fasta.gz](ftp://ftp.flybase.net/genomes/dmel/dmel_r5.57_FB2014_03/fasta/dmel-all-gene-r5.57.fasta.gz).

The annotated gff file was downloaded from:

[ftp://ftp.flybase.net/genomes/dmel/dmel_r5.57_FB2014_03/gff/dmel-all-r5.57.gff.gz](ftp://ftp.flybase.net/genomes/dmel/dmel_r5.57_FB2014_03/gff/dmel-all-r5.57.gff.gz).

Transposable element (TE) features were extracted and converted to bed file using:

```
awk'$2=="FlyBase"&&$3=="transposable_element" {print $0}' <dmel-all-r5.57.gff> >
<TE.gtf>
awk'{print $1"\t"$4"\t"$5"\t"$3"_"NR}' <TE.gtf> > <TE.bed>
```

Then, the pipeline was executed as:

```
assembled_feature_pipeline.sh -a <asm.fasta> -r <reference.fasta> -f <TE.bed>
```

After running the scripts, the final output file, called results/FINAL.REPORT, was generated and used to identify TE with ⩾100% Pct_length and corresponding Pct_ident. The repeat familes, *roo* and *juan*, were extracted from the FINAL.REPORT file. The annotated region from *roo* and *juan* families can be extracted from dmel-all-r5.57.gff. The results are shown in Supplementary Table 9.

### Supplementary Note 12: Analysis of telomere assembly

LRs provide considerable advantages in reconstructing the repetitive heterochromatic regions of eukaryotic chromosomes. Telomeres play important roles in chromosome replication of all eukaryotic genomes. Nanopore LR sequencing presents distinct advantages in telomere assembly. To validate the effectiveness of using Nanopore data, we evaluated long-read sequencing in reconstruction of heterochromatic sequences in telomeric regions of *S. cerevisiae*.

The *S. cerevisiae* S288C other features database was downloaded from http://downloads.yeastgenome.org/sequence/S288C_reference/other_features/other_features_genomic.fasta.gz. We mapped selected *S. cerevisiae* telomeric repeats to *S. cerevisiae* W303 assemblies generated using Canu, Canu+Smartdenovo, Smartdenove, miniasm, wtdbg2, Flye and NECAT.

The features were aligned to the assembly using the following scripts:

```
nucmer --maxmatch<asm.fasta><features.fasta>

show-coords -lrcTHout.delta |sort -nk12 |awk'{if ($7> 85 && $11> 50) print $0}'
| grep TEL | sort -rnk8 >tels.coords
```

The contigs containing telomeric features within 1 kbp of contig ends were then identified. The results are shown in Supplementary Table 10.

### Supplementary Note 13: Validation of *H. sapiens* NA12878

To validate the performances of NECAT and Canu, each polished assembly was aligned to reference genome hg38 with MUMmer (v4.0)[7], after which tiling figures were generated (**Supplementary Figure 12)**. The genome was polished four times using Nanopore data with Racon[6] (v1.3.1) and minimap2[23] (v2.10-r761), after which the final genome was generated using the following code:

```
minimap2 -x map-ont -t $NPROC $DRAFT reads.fastq > ONTmin_IT0.paf
time racon -m 8 -x -6 -g -8 -w 500 -t $NPROC reads.fastq ONTmin_IT0.paf $DRAFT > ONTmin_IT1.fasta
minimap2 -x map-ont -t $NPROC ONTmin_IT1.fasta reads.fastq > ONTmin_IT1.paf
time racon -m 8 -x -6 -g -8 -w 500 -t $NPROC reads.fastq ONTmin_IT1.paf ONTmin_IT1.fasta > ONTmin_IT2.fasta
minimap2 -x map-ont -t $NPROC ONTmin_IT2.fasta reads.fastq > ONTmin_IT2.paf
time racon -m 8 -x -6 -g -8 -w 500 -t $NPROC reads.fastq ONTmin_IT2.paf ONTmin_IT2.fasta > ONTmin_IT3.fasta
minimap2 -x map-ont -t $NPROC ONTmin_IT3.fasta reads.fastq > ONTmin_IT3.paf
time racon -m 8 -x -6 -g -8 -w 500 -t $NPROC reads.fastq ONTmin_IT3.paf ONTmin_IT3.fasta > ONTmin_IT4.fasta
```

The custom scripts, used to convert the output into a format accepted by ColoredChromesomes.pl (http://sourceforge.net/projects/cchrom/), are shown below:

```
python makeMappings.py asm_refhg38.1coords 10000 > asm.tiling
perl convertToChr.pl human.chr.map asm.tiling human.lanes human.chrPos > asm.cfg
perl coloredChromosomes.pl --chromosomeSpec asm.cfg -o asm.ps
ps2pdf asm.ps
```

Because MUMmer was set to use a unique anchor matching option to accelerate the alignment, some repetitive sequences remained unaligned. To avoid displaying these regions as gaps in tiling, the conversion script chained together consecutive alignments from the same contig if alignment gap in the reference was less than 10,000 bp. Thus, breaks in the resulting tiling occurred whenever a contig switch occurred, or if there was a >10,000 bp gap between two alignments of the same contig. The entire process of alignment and figure generation can be reproduced using the scripts available on the MHAP home page[20] (assuming that Perl, Python, and MUMmer[7] are placed in the correct path), and by running the script shown below; this generates a figure designated as asm.pdf (**Supplementary Figure 14**).

```
sh makeHuman.sh ref.fasta asm.fasta
```
