## Supplementary Tables for "Fast and accurate assembly of Nanopore reads via progressive error correction and adaptive read selection"

**Supplementary Table 1. Detail information of the six datasets used in this study.**

| Datasets | Data resource of LRs | SRA ID of NGS | Reference |
|---|---|---|---|
| *E. coli* | Ecoli.fasta<br>ecoli.fast5.tgz | SRR072235 | K-12 substr. MG1655 |
| *S. cerevisiae* | yeast.fastq<br>yeast.fast5.tar.gz | SRR5244182 | *S. cerevisiae* S288c |
| *D. melanogaster* | SRX3676783<br>Dro1.fast5.tar.gz<br>Dro2.fast5.tar.gz | SRR6702604 | *D. melanogaster* v6 |
| *A. thaliana* | ERR2173373<br>Ara.bam | ERR2173372 | *A. thaliana* v5 |
| *C. reinhardtii* | chl.fastq.gz<br>chl.fast5.tar.gz | SRR1734612 | *C. reinhardtii* v5.5 |
| *O. sativa* | rice.fastq.gz | — | O. sativa v4.0 |
| *S. pennellii* | tomatodata | — | S. pennellii |
| *H. sapiens*<br>(NA12878) | rel_3_4 data<br>fast5 data | ERR194147 | Hg38 |

LRs: long reads; NGS: Next generation sequencing

**Supplementary Table 2. Statistical information of the eight datasets used in this study.**

| Datasets | Base size | Coverage | LR Count | N25 | N50 | N75 | Mean |
|---|---|---|---|---|---|---|---|
| *E. coli* | 1,481,822,528 | 322X | 164,472 | 25,243 | 14,891 | 8,074 | 9,010 |
| *S. cerevisiae* | 7,354,232,165 | 612X | 1,814,834 | 27,799 | 13,354 | 4,108 | 4,052 |
| *D. melanogaster* | 9,064,470,438 | 66X | 1,327,569 | 18,919 | 11,853 | 6,495 | 6,828 |
| *A. thaliana* | 3,421,779,258 | 27X | 300,071 | 30,160 | 20,127 | 11,544 | 11,403 |
| *C. reinhardtii* | 16,124,079,751 | 134X | 1,455,141 | 45,167 | 27,507 | 14,469 | 11,080 |
| *O. sativa* | 67,710,180,969 | 183X | 2,696,991 | 46,050 | 33,120 | 21,676 | 25,106 |
| *S. pennellii* | 141,704,928,841 | 160X | 11,967,377 | 21,210 | 16,611 | 12,552 | 11,841 |
| *H. sapiens* (NA12878) | 114,380,310,980 | 38X | 15,599,457 | 22,232 | 12,196 | 7,209 | 7,332 |

HERS: high error subsequences with high sequencing error rates > 50% in 500bp subsequence.

**Supplementary Table 3. Statistical information of sequencing error rate for eight datasets.**

| **Datasets** | **The mean error rate of raw reads** | **Percentage of reads with HERS and length>10kb** |
|---|---|---|
| *E.coli* | 17.80% | 3.50% |
| *S. cerevisiae* | 12.00% | 5.20% |
| *A. thaliana* | 20.10% | 23.2% |
| *D. melanogaster* | 16.20% | 6.90% |
| *C. reinhardtii* | 15.00% | 10.90% |
| *O. sativa* | 15.60% | 8.10% |
| *S. pennellii* | 18.49% | 8.05% |
| Human(NA12878) | 19.50% | 22.50% |

HERS: high error subsequences with high sequencing error rates > 50% in 1000bp subsequence.

**Supplementary Table 4. Comparison of the accuracy of the five datasets used in this study.**

| Species | Error rate | Percentage | | | |
|---|---|---|---|---|---|
| | | Raw | Canu | CN1 | CN2 |
| *A. thaliana* | 1% | 0.10% | 0.08% | 0.13% | 0.29% |
| | 2% | 0.19% | 0.32% | 0.82% | 3.51% |
| | 3% | 0.33% | 0.80% | 3.07% | 15.08% |
| | 4% | 0.42% | 1.64% | 8.86% | 17.27% |
| | 5% | 0.53% | 5.25% | 12.79% | 9.70% |
| | 6% | 0.69% | 10.27% | 9.60% | 6.62% |
| | 7% | 0.99% | 10.13% | 7.06% | 5.28% |
| | 8% | 1.29% | 8.25% | 5.83% | 4.51% |
| | 9% | 1.36% | 6.72% | 4.91% | 3.98% |
| | 10% | 1.47% | 5.73% | 4.37% | 3.78% |
| | 10-15% | 21.93% | 19.98% | 17.24% | 12.89% |
| | 15-20% | 23.94% | 13.53% | 11.63% | 7.01% |
| | 20-15% | 19.40% | 8.44% | 5.88% | 3.90% |
| | 25-30% | 14.37% | 4.27% | 3.34% | 2.65% |
| | 30-100% | 13.00% | 4.60% | 4.47% | 3.52% |
| *E. coli* | 1% | 0.00% | 0.00% | 0.00% | 0.21% |
| | 2% | 0.00% | 0.02% | 0.57% | 38.03% |
| | 3% | 0.00% | 0.12% | 13.41% | 54.65% |
| | 4% | 0.00% | 6.49% | 39.09% | 5.51% |
| | 5% | 0.01% | 13.82% | 27.44% | 0.94% |
| | 6% | 0.05% | 14.93% | 10.95% | 0.30% |
| | 7% | 0.12% | 17.18% | 4.26% | 0.12% |
| | 8% | 0.26% | 16.45% | 1.83% | 0.06% |
| | 9% | 0.52% | 12.85% | 0.90% | 0.03% |
| | 10% | 1.10% | 8.23% | 0.46% | 0.03% |
| | 10-15% | 30.91% | 9.36% | 0.66% | 0.05% |
| | 15-20% | 37.18% | 0.44% | 0.19% | 0.02% |
| | 20-15% | 20.07% | 0.07% | 0.09% | 0.01% |
| | 25-30% | 8.53% | 0.03% | 0.06% | 0.01% |
| | 30-100% | 1.23% | 0.01% | 0.10% | 0.01% |
| *S. cerevisiae* | 1% | 0.01% | 8.93% | 22.94% | 73.24% |
| | 2% | 0.03% | 31.27% | 32.96% | 13.85% |
| | 3% | 0.16% | 24.28% | 17.77% | 4.26% |
| | 4% | 0.49% | 14.81% | 9.57% | 2.37% |
| | 5% | 0.92% | 8.01% | 4.85% | 1.32% |
| | 6% | 1.71% | 4.28% | 2.64% | 0.76% |
| | 7% | 4.74% | 2.42% | 1.58% | 0.50% |
| | 8% | 9.05% | 1.53% | 1.10% | 0.38% |
| | 9% | 11.09% | 1.02% | 0.84% | 0.34% |
| | 10% | 10.47% | 0.67% | 0.65% | 0.20% |
| | 10-15% | 35.62% | 1.60% | 2.22% | 1.02% |
| | 15-20% | 16.69% | 0.53% | 1.15% | 0.63% |
| | 20-15% | 6.44% | 0.25% | 0.60% | 0.40% |
| | 25-30% | 1.41% | 0.17% | 0.37% | 0.25% |
| | 30-100% | 1.16% | 0.23% | 0.75% | 0.49% |
| *D. melanogaster* | 1% | 0.41% | 1.40% | 4.62% | 39.13% |
| | 2% | 0.44% | 20.23% | 24.01% | 24.44% |
| | 3% | 0.43% | 20.70% | 20.10% | 4.65% |
| | 4% | 0.47% | 9.79% | 10.30% | 2.03% |
| | 5% | 0.55% | 5.45% | 5.15% | 1.78% |
| | 6% | 0.73% | 3.98% | 3.30% | 1.84% |
| | 7% | 1.35% | 3.01% | 2.68% | 1.43% |

| | | | | | |
|---|---|---|---|---|---|
| | 8% | 2.83% | 2.29% | 2.20% | 1.03% |
| | 9% | 4.91% | 1.84% | 1.69% | 1.34% |
| | 10% | 6.62% | 1.91% | 1.34% | 2.51% |
| | 10-15% | 32.94% | 8.34% | 7.01% | 9.35% |
| | 15-20% | 21.64% | 8.52% | 8.18% | 5.16% |
| | 20-15% | 13.97% | 6.89% | 5.24% | 2.89% |
| | 25-30% | 7.91% | 3.34% | 2.46% | 1.44% |
| | 30-100% | 4.79% | 2.31% | 1.71% | 0.99% |
| *C. reinhardtii* | 1% | 0.40% | 0.10% | 0.96% | 36.91% |
| | 2% | 0.24% | 9.99% | 17.07% | 42.08% |
| | 3% | 0.18% | 35.85% | 33.06% | 10.98% |
| | 4% | 0.16% | 20.99% | 21.11% | 3.56% |
| | 5% | 0.18% | 9.12% | 9.93% | 1.65% |
| | 6% | 0.39% | 4.87% | 4.84% | 0.94% |
| | 7% | 0.69% | 3.33% | 2.76% | 0.59% |
| | 8% | 0.75% | 2.32% | 1.78% | 0.42% |
| | 9% | 1.11% | 1.73% | 1.24% | 0.33% |
| | 10% | 3.28% | 1.27% | 0.91% | 0.25% |
| | 10-15% | 61.31% | 3.64% | 2.41% | 0.75% |
| | 15-20% | 17.43% | 1.91% | 1.14% | 0.42% |
| | 20-15% | 6.35% | 1.52% | 0.77% | 0.39% |
| | 25-30% | 3.16% | 1.27% | 0.66% | 0.27% |
| | 30-100% | 4.39% | 2.10% | 1.35% | 0.48% |
| *O. sativa* | 1% | 0.05% | 0.13% | 0.25% | 4.85% |
| | 2% | 0.06% | 1.63% | 5.32% | 24.66% |
| | 3% | 0.09% | 11.43% | 14.88% | 20.55% |
| | 4% | 0.12% | 15.56% | 15.99% | 14.56% |
| | 5% | 0.17% | 15.67% | 15.05% | 10.00% |
| | 6% | 0.27% | 12.70% | 12.18% | 6.73% |
| | 7% | 0.43% | 9.10% | 9.15% | 4.28% |
| | 8% | 0.65% | 6.30% | 6.61% | 2.49% |
| | 9% | 1.14% | 4.19% | 4.61% | 1.33% |
| | 10% | 2.47% | 2.62% | 2.96% | 0.77% |
| | 10-15% | 42.41% | 5.63% | 5.16% | 3.63% |
| | 15-20% | 42.39% | 5.85% | 3.47% | 3.25% |
| | 20-15% | 5.46% | 5.30% | 2.82% | 1.69% |
| | 25-30% | 2.75% | 2.57% | 0.88% | 0.64% |
| | 30-100% | 1.56% | 1.33% | 0.66% | 0.58% |
| *S. pennellii* | 1% | 0.18% | 0.10% | 0.45% | 3.23% |
| | 2% | 0.18% | 1.92% | 3.83% | 17.68% |
| | 3% | 0.27% | 7.76% | 8.96% | 20.36% |
| | 4% | 0.43% | 12.46% | 12.90% | 13.31% |
| | 5% | 0.64% | 11.80% | 12.63% | 8.46% |
| | 6% | 0.90% | 9.45% | 10.08% | 5.86% |
| | 7% | 1.25% | 7.37% | 7.71% | 4.17% |
| | 8% | 1.80% | 5.55% | 5.93% | 3.14% |
| | 9% | 2.52% | 4.15% | 4.53% | 2.60% |
| | 10% | 3.40% | 3.16% | 3.46% | 2.46% |
| | 10-15% | 26.81% | 15.57% | 12.25% | 7.85% |
| | 15-20% | 25.30% | 11.24% | 6.90% | 4.17% |
| | 20-15% | 16.76% | 5.23% | 3.89% | 2.76% |
| | 25-30% | 10.10% | 2.18% | 2.41% | 2.05% |
| | 30-100% | 9.48% | 2.07% | 4.09% | 1.89% |

### Supplementary Table 7. Comparison with assemble-then-correct assemblers

| Genome | Pipeline | Assembly Size | Contig | NG50 (AP) | ctg/ chr | Total time |
|---|---|---|---|---|---|---|
| *E. coli* | Ref. | 4641652 | 1 | 4,641,652(100%) | 1 | — |
| | Miniasm | 4407447 | 1 | 4,407,447(95%) | 1 | 2.9 |
| | Smartdenovo | 4631621 | 1 | 4,631,621(100%) | 1 | 40 |
| | Wtdbg2 | 4494759 | 1 | 4,494,759(97%) | 1 | 0.8 |
| | Flye | 4622475 | 1 | 4,622,475(100%) | 1 | 630.4 |
| | NECAT | 4594537 | 1 | 4,594,537(99%) | 1 | 2.8 |
| *S. cerevisiae* | S228C | 12157105 | 17 | 924,431(100%) | 1 | — |
| | Miniasm | 13005314 | 33 | 820,023(89%) | 2 | 63.5 |
| | Smartdenovo | 12428603 | 20 | 936,846(101%) | 1 | 97.1 |
| | Wtdbg2 | 12054756 | 22 | 791,587(86%) | 1 | 6.3 |
| | Flye | 12327296 | 26 | 943,486(102%) | 2 | 197.8 |
| | NECAT | 12341147 | 19 | 936,684(101%) | 1 | 9.3 |
| *A. thaliana* | TAIR10 | 119668634 | 7 | 23,459,830(100%) | 1 | — |
| | Miniasm | 112043421 | 71 | 11,303,164(48%) | 10 | 5.87 |
| | Smartdenovo | 116434478 | 127 | 3,676,347(16%) | 18 | 78.4 |
| | Wtdbg2 | 115323989 | 349 | 9,840,213(42%) | 50 | 14.4 |
| | Flye | 126613812 | 154 | 12,042,914(51%) | 22 | 59.4 |
| | NECAT | 122855840 | 136 | 11,157,362(48%) | 19 | 47.9 |
| *D. melanogaster* | dm5 | 143726002 | 1870 | 25,286,936(100%) | 234 | — |
| | Miniasm | 139880718 | 443 | 1,349,941(5%) | 55 | 36.3 |
| | Smartdenovo | 138142528 | 238 | 4,479,808(18%) | 30 | 182.4 |
| | Wtdbg2 | 138929862 | 872 | 6,633,247(26%) | 109 | 26.0 |
| | Flye | 139860664 | 593 | 11,924,703(47%) | 74 | 127.9 |
| | NECAT | 142774092 | 277 | 18,072,166(71%) | 35 | 70.4 |
| *C. reinhardtii* | Ref. v3.0 | 111098438 | 53 | 7,783,580(100%) | 3 | — |
| | Miniasm | 122181355 | 215 | 2,388,992(31%) | 13 | 118.9 |
| | Smartdenovo | 112895049 | 83 | 3,370,315(43%) | 5 | 1365.3 |
| | Wtdbg2 | 115667394 | 344 | 4,289,786(55%) | 22 | 35.4 |
| | Flye | 112901001 | 65 | 6,572,682(84%) | 4 | 185.8 |
| | NECAT | 113388358 | 54 | 6,168,830(79%) | 3 | 101.8 |
| *O. sativa* | Ref.v4.0 | 382778125 | 15 | 30,828,668(100%) | 1 | — |
| | Miniasm | 383376696 | 240 | 9,511,261(31%) | 16 | 682.0 |
| | Smartdenovo | 379412051 | 352 | 1,888,744(6%) | 23 | 3564.9 |
| | Wtdbg2 | 394595916 | 2554 | 2,432,307(8%) | 170 | 154.3 |
| | Flye | 380658776 | 249 | 3,552,487(12%) | 17 | 937.3 |
| | NECAT | 373120604 | 120 | 9,650,275(31%) | 8 | 517.2 |
| *S. pennellii* | Ref.SL.3.0 | 915596307 | 899 | 2,521,711(100%) | 69 | — |
| | Miniasm | 977783143 | 2704 | 1,903,403(75%) | 208 | — |
| | Smartdenovo | 955307836 | 1901 | 1,108,002(44%) | 146 | — |
| | Wtdbg2 | 934260260 | 4986 | 1,227,952(49%) | 384 | 439.0 |
| | Flye | 1025983422 | 3180 | 1,971,110(78%) | 245 | 4422.8 |
| | NECAT | 991792915 | 1344 | 4,801,589(190%) | 103 | 2540.3 |

Assembly size is the total number of base pairs in all contigs generated by assemblers. NG50 indicates that 50% of reference genome size was contained in contigs having length ≥N. Assembly performance (AP) is defined as obtained contig NG50 divided by NG50 of reference assembly. The genome sizes of *E. coli, S. cerevisiae* W303, *A. thaliana* Col-0, *D. melanogaster* ISO1, *C. reinhardtii, O. sativa* and *S. pennellii* are 4,641,652, 12,157,105, 119,668,634, 143,726,002, 111,098,438, 382,778,125, and 915,596,307, respectively. Ctg/Chr is the average number of contigs per chromosome in the assembly. All the pipelines were tested on the same computer with 2.0 GHz CPU and 3T GB RAM of memory. For the first six datasets, we ran all the pipelines on our computer with 32 threads; the total computational time are recorded. For *S. pennellii* dataset, we ran the pipelines on our computer with 64 threads, and total computational time are recorded. The assembly results of Miniasm and Smartdenove on the dataset *S. pennellii* were from https://www.plabipd.de/portal/solanum-pennellii.

**Supplementary Table 8. SNP and INDEL statistics between assembly genome and reference genome.**

| Species | Software | SNPs | Indels | | Cover (%) |
|---|---|---|---|---|---|
| | | | <=10bp | >10bp | |
| *E. coli* | Canu | 9384 | 0 | 19 | 99.34 |
| | Canu1.8 | 13010 | 19 | 45 | 98.02 |
| | Canu_smartdenovo | 9170 | 0 | 4 | 99.39 |
| | Smartdenovo | 8663 | 0 | 6 | 99.31 |
| | miniasm | 19355 | 0 | 3 | 98.59 |
| | wtdbg2 | 7926 | 0 | 4 | 99.39 |
| | Flye | 8590 | 0 | 3 | 99.42 |
| | NECAT | 10248 | 0 | 11 | 99.37 |
| *S. cerevisiae* | Canu | 9259 | 5 | 36 | 99.78 |
| | Canu1.8 | 9297 | 7 | 33 | 99.88 |
| | Canu_smartdenovo | 9010 | 4 | 30 | 99.91 |
| | Smartdenovo | 9699 | 8 | 41 | 99.65 |
| | miniasm | 9581 | 12 | 37 | 99.56 |
| | wtdbg2 | 9243 | 7 | 31 | 99.82 |
| | Flye | 9199 | 8 | 31 | 99.92 |
| | NECAT | 9142 | 4 | 38 | 99.92 |
| *D. melanogaster* | Canu | 24724 | 2 | 299 | 99.03 |
| | Canu1.8 | 24811 | 22 | 266 | 98.96 |
| | Canu_smartdenovo | 31359 | 9 | 281 | 99.16 |
| | Smartdenovo | 40735 | 5 | 251 | 98.92 |
| | miniasm | 43817 | 3 | 217 | 99.07 |
| | wtdbg2 | 27297 | 24 | 215 | 98.87 |
| | Flye | 26494 | 20 | 195 | 99.03 |
| | NECAT | 29113 | 10 | 345 | 99.30 |
| *A. thaliana* | Canu | 458066 | 38 | 3312 | 98.98 |
| | Canu1.8 | 457782 | 456 | 2880 | 98.98 |
| | Canu_smartdenovo | 462818 | 42 | 3413 | 99.32 |
| | Smartdenovo | 461263 | 37 | 3372 | 99.02 |
| | miniasm | 462697 | 43 | 3406 | 99.24 |
| | wtdbg2 | 457763 | 458 | 2976 | 99.19 |
| | Flye | 463692 | 465 | 2988 | 99.34 |
| | NECAT | 463859 | 45 | 3427 | 99.31 |
| *C. reinhardtii* | Canu | 40645 | 15 | 1902 | 99.26 |
| | Canu1.8 | 39984 | 324 | 1580 | 99.40 |
| | Canu_smartdenovo | 39832 | 13 | 1630 | 99.60 |
| | Smartdenovo | 47750q | 12 | 1655 | 99.40 |
| | miniasm | 42732 | 12 | 1605 | 99.48 |
| | wtdbg2 | 48849 | 283 | 1372 | 99.21 |
| | Flye | 39996 | 302 | 1388 | 99.69 |
| | NECAT | 45218 | 11 | 1812 | 99.54 |

**Supplementary Table 9. Number of TEs in Flybase.**

| Method | Contain in a contig | | *roo* | | *Juan* | |
|---|---|---|---|---|---|---|
| | Total | Perfect | Total | Perfect | Total | Perfect |
| Canu | 5304 | 3970 | 131 | 95 | 11 | 11 |
| Canu+Smartdenovo | 5292 | 3916 | 132 | 115 | 11 | 11 |
| Miniasm | 5244 | 3924 | 132 | 77 | 9 | 9 |
| Smrtdenovo | 5312 | 3998 | 135 | 106 | 11 | 11 |
| Flye | 5268 | 3840 | 131 | 93 | 11 | 11 |
| Wtdbg2 | 5156 | 3831 | 130 | 95 | 11 | 11 |
| NECAT | 5,304 | 4001 | 134 | 118 | 11 | 11 |

Total and Perfect refer to all the identified TE numbers and the number of TEs with more than 99% Pct_ident from the final report.

**Supplementary Table 10. Chromosome number identified based on the alignment of telomeric repeats**

| Method | All | Pair_end telomere | | Single_end telomere |
|---|---|---|---|---|
| | | Identified in a single contigs | Identified in two contigs | |
| Canu | 16 | 13 | 1 | 2 |
| Canu+Smartdenovo | 16 | 14 | 1 | 1 |
| Miniasm | 16 | 13 | 1 | 2 |
| Smrtdenovo | 16 | 14 | 1 | 1 |
| Flye | 16 | 14 | 1 | 1 |
| Wtdbg2 | 16 | 3 | 8 | 5 |
| NECAT | 16 | 14 | 1 | 1 |

"All" indicates the total identified number of chromosome, "identified in a single contig" is the number of chromosomes in which the telomeric repeats are mapped on both the left and right ends in a single contig.
