## Supplementary Figures for "Fast and accurate assembly of Nanopore reads via progressive error correction and adaptive read selection"

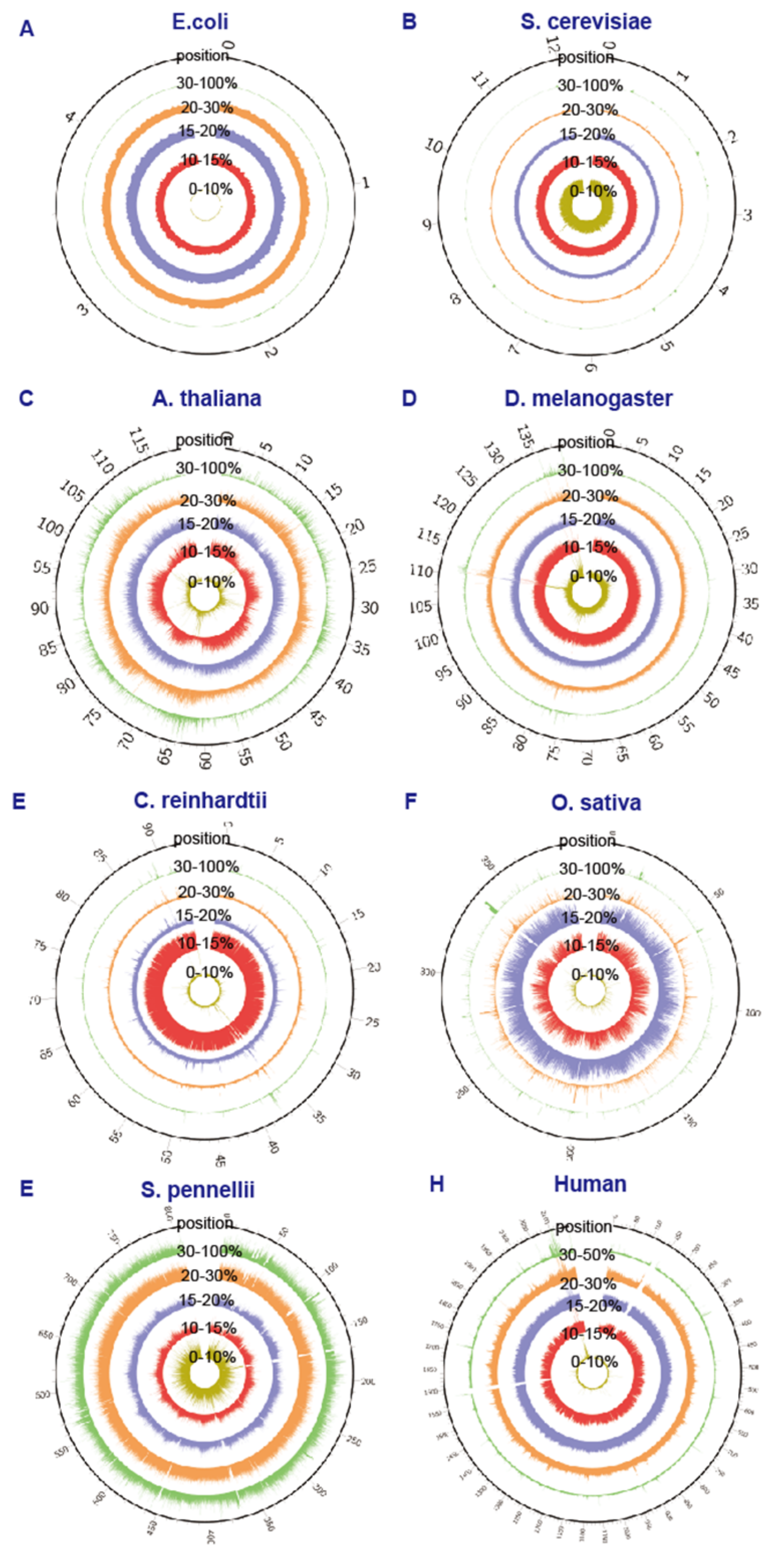


Supplementary Figure 1

Sequencing error distribution for aligned Nanopore raw long reads on different reference-genome positions in the nanopore datasets. (I: genome position; II: percentage of reads with 30-100% sequencing error rate; III: 25-30%; IV: 20-25%; V: 15-20%; VI: 10-15%; VII: 0-10%).

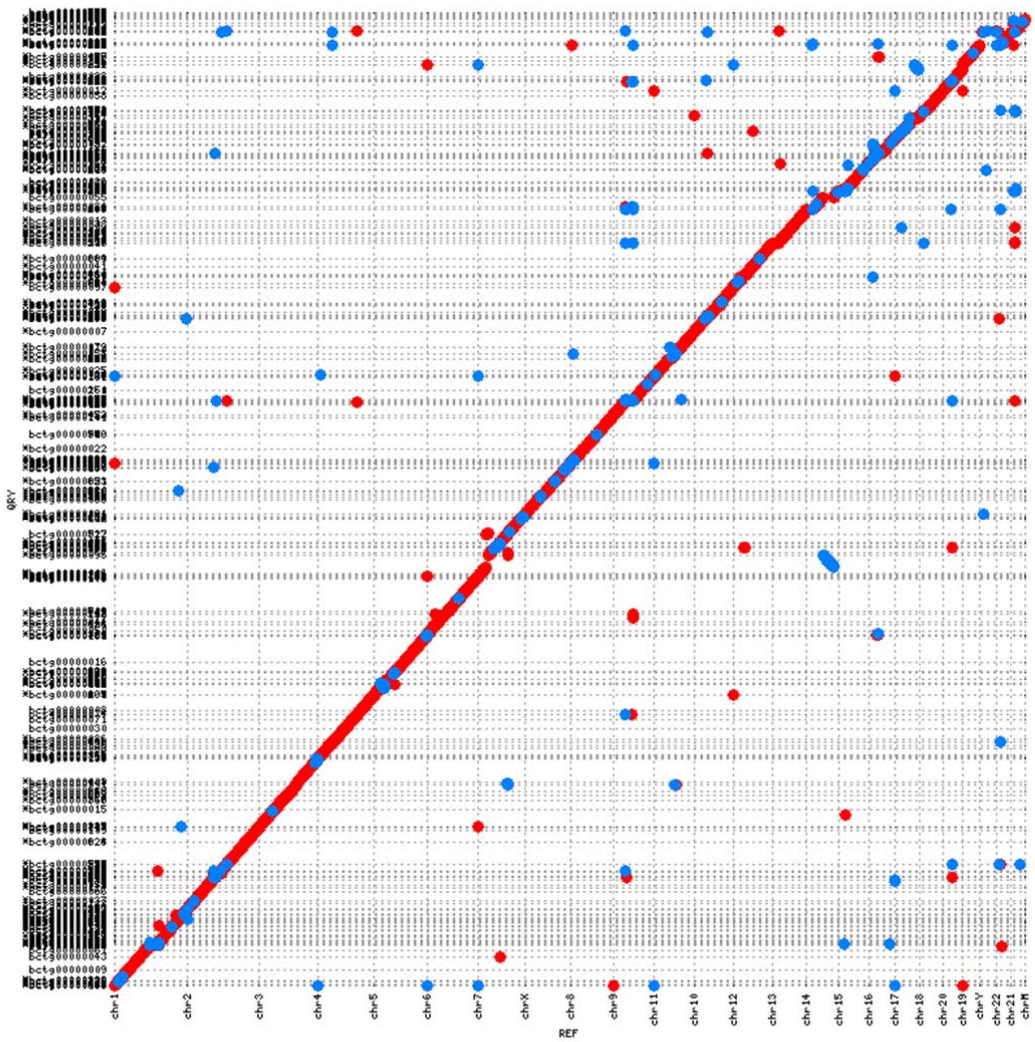


Supplementary Figure 2

Mummerplot of new assembled WERI Nanopore contigs and hg38 reference genome. An alignment dotplot shows the relationship between the contig assembled using Nanopore (y-axis) and GRCh38 reference genome (x-axis). Contig and chromosome boundaries are displayed as dotted lines (horizontal and vertical, respectively).

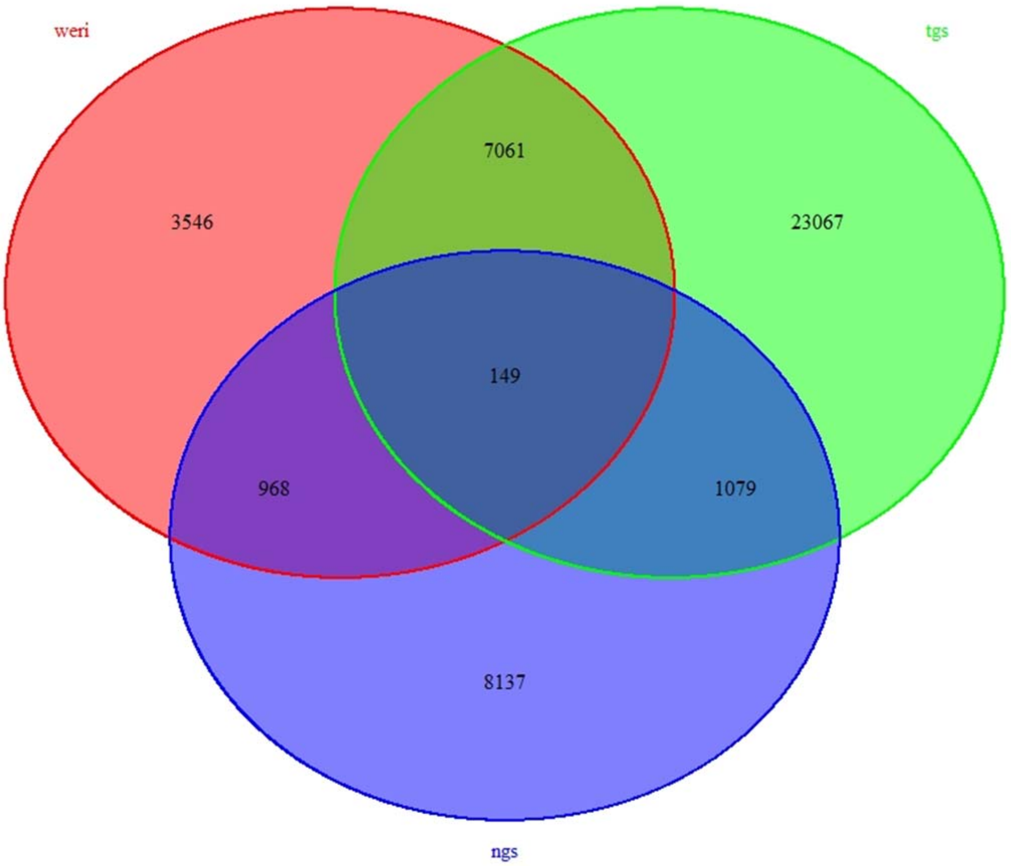


Supplementary Figure 3

The number of identified SVs detected with WERI, TGS, and NGS.

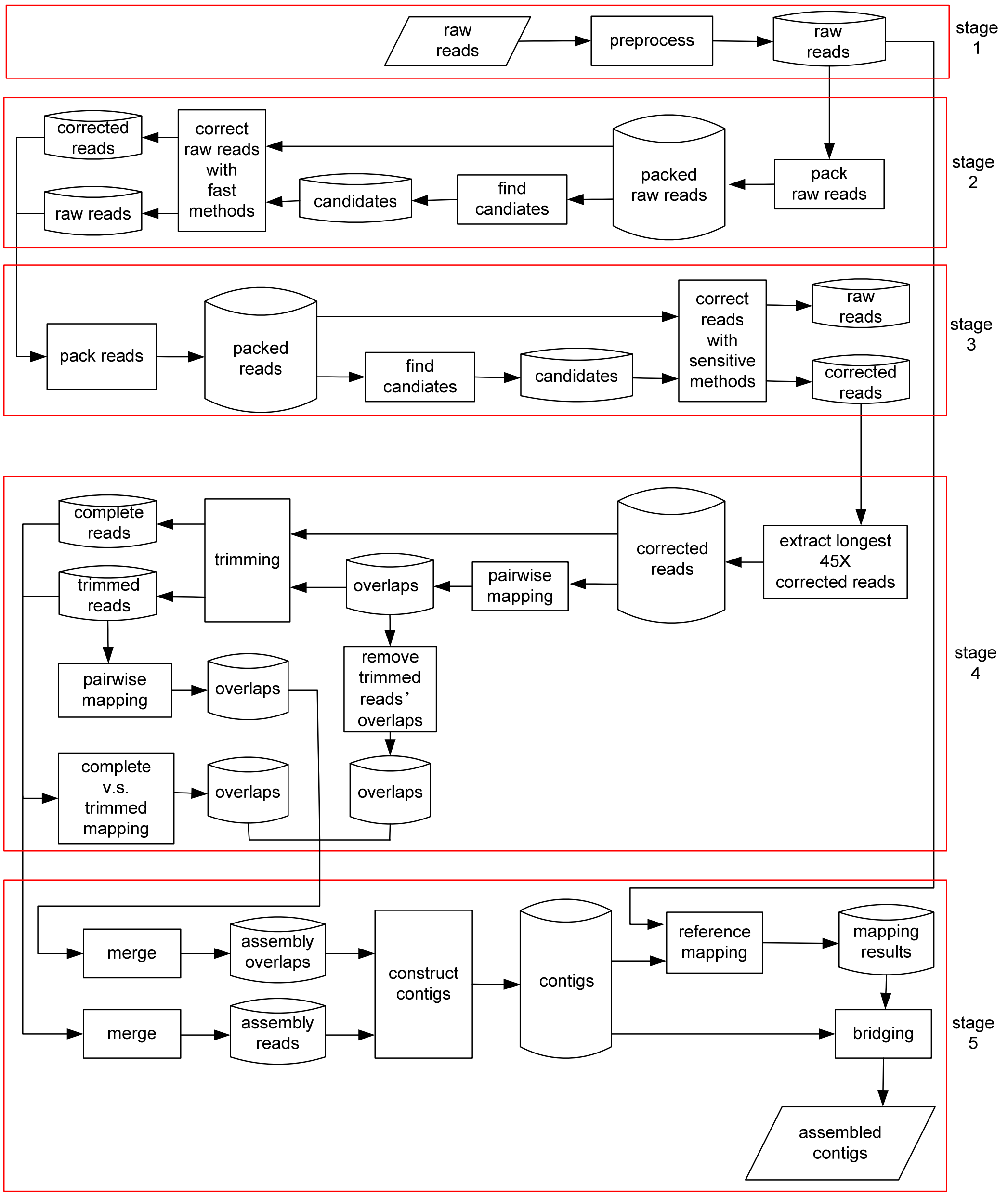


Supplementary Figure 4

NECAT architecture. Stage 1: preprocess; Stage 2: step one of correction; Stage 3: step two of correction; Stage 4: trimming; Stage 5: assembly.

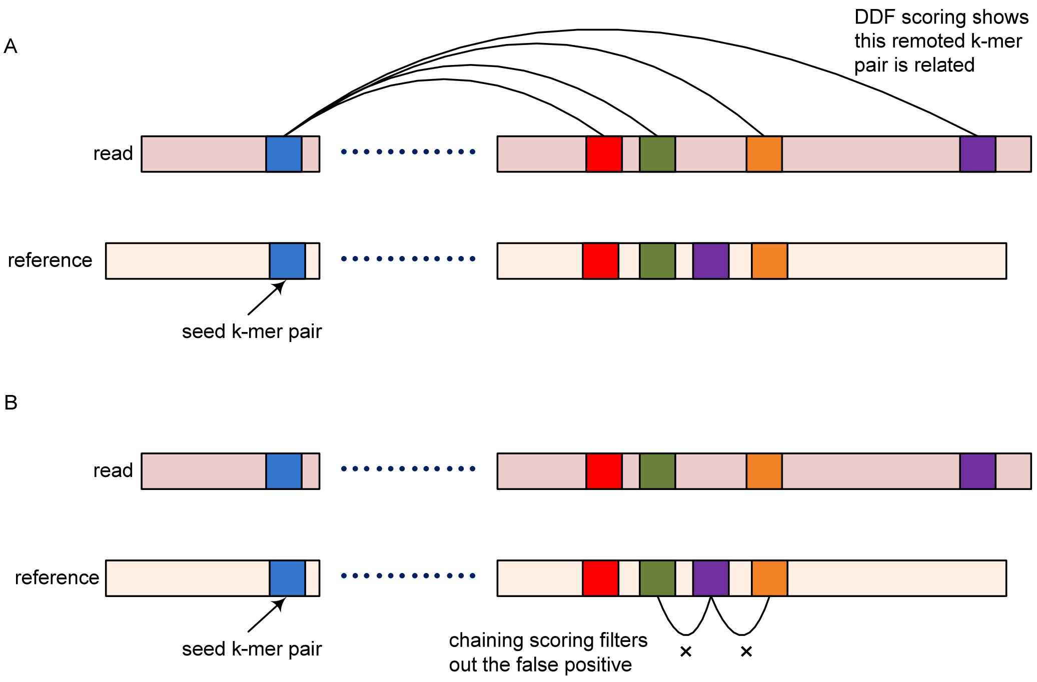


Supplementary Figure 5

Removing false positives with chaining technique. (A): The candidate k-mer pair (blue) and its four remote related k-mer pairs detected by DDF scoring. (B)Chaining is used to remove false positives (purple) by examining positions with adjacent k-mer pairs.

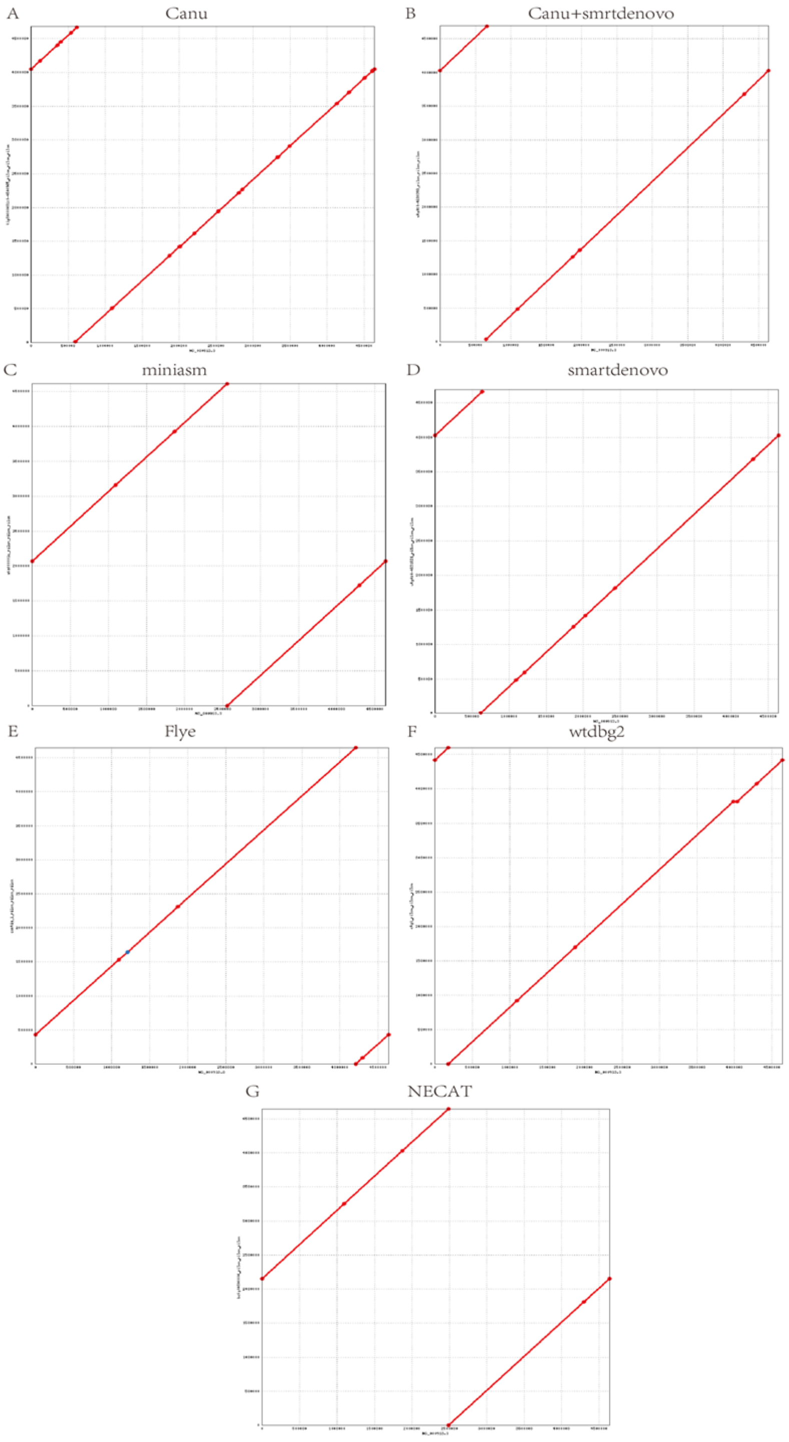


Supplementary Figure 6

Mummerplot of assembled contig and *E. coli* reference genome. An alignment dotplot shows the relationship between the assembled contig of *E. coli* K12 (y-axis) and *E. coli* K12 reference genome (x-axis). The assembled single contig was mapped onto the reference genome and covered the entire genome. The assembled contig was arbitrarily shifted because the *E. coli* chromosome is circular; this does not represent assembly error.

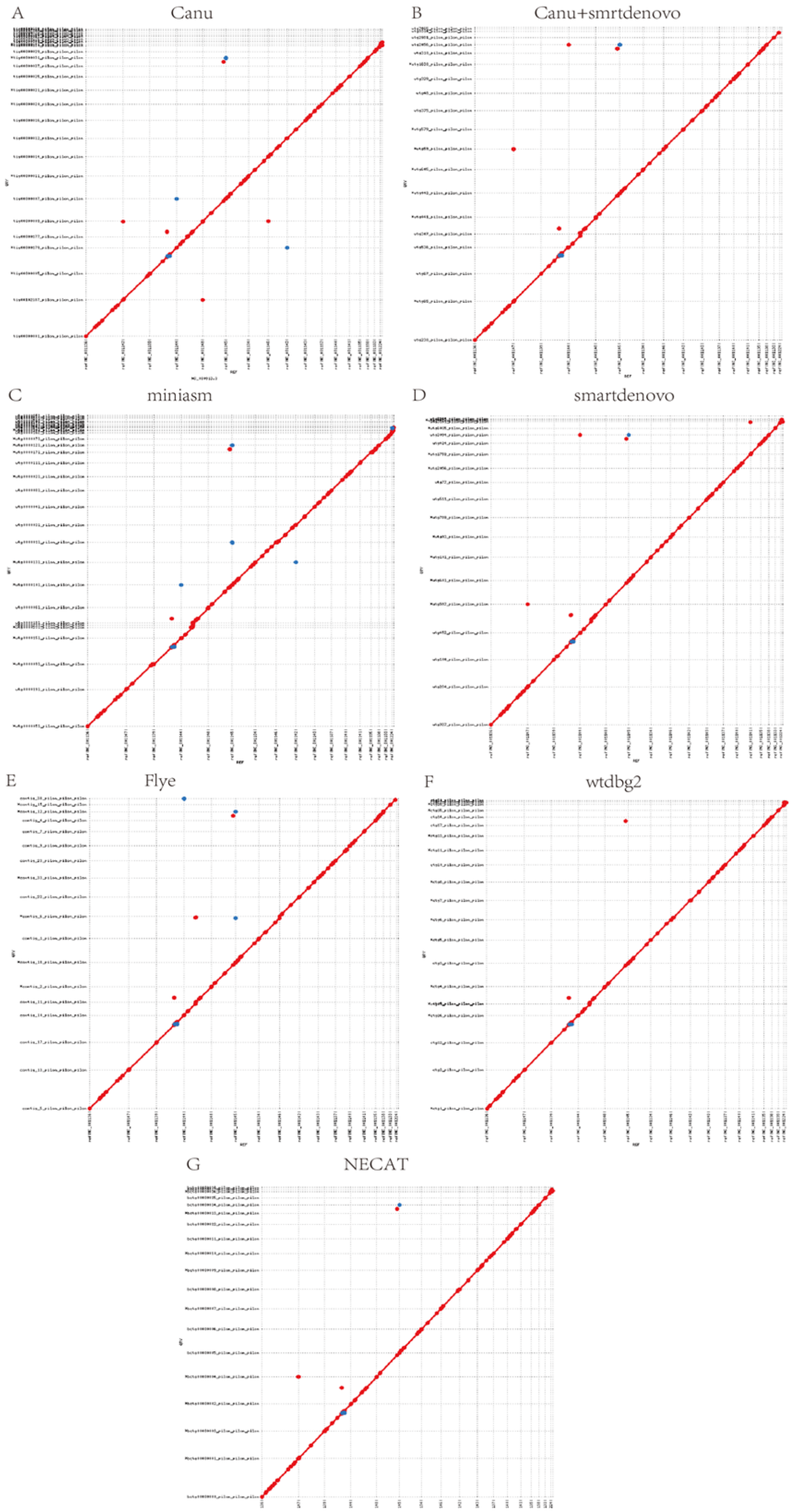


Supplementary Figure 7

Mummerplot of the assembled contig and *S. cerevisiae* reference genome.

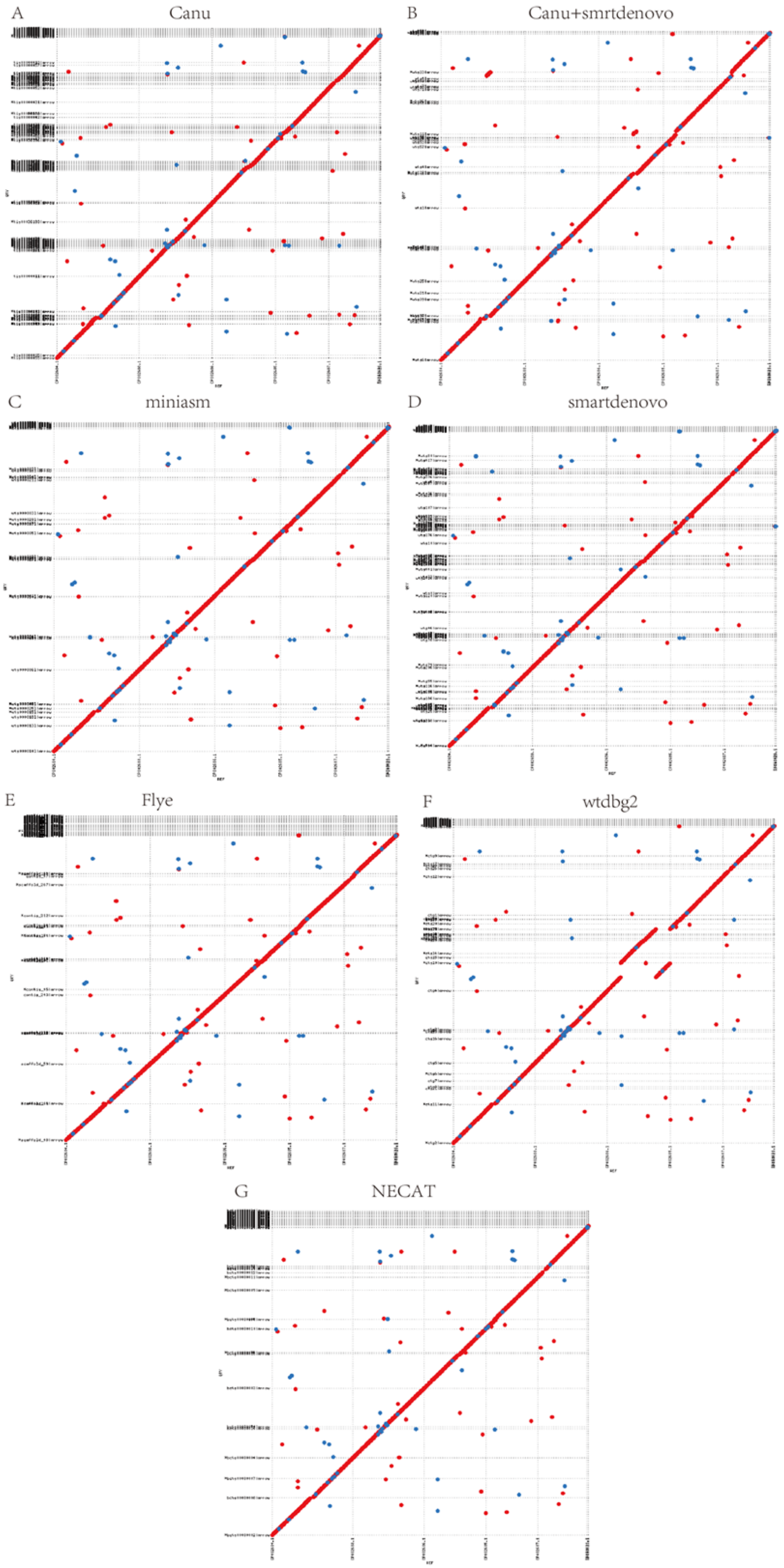


Supplementary Figure 8

Mummerplot of the assembled contig and *A. thaliana* reference genome.

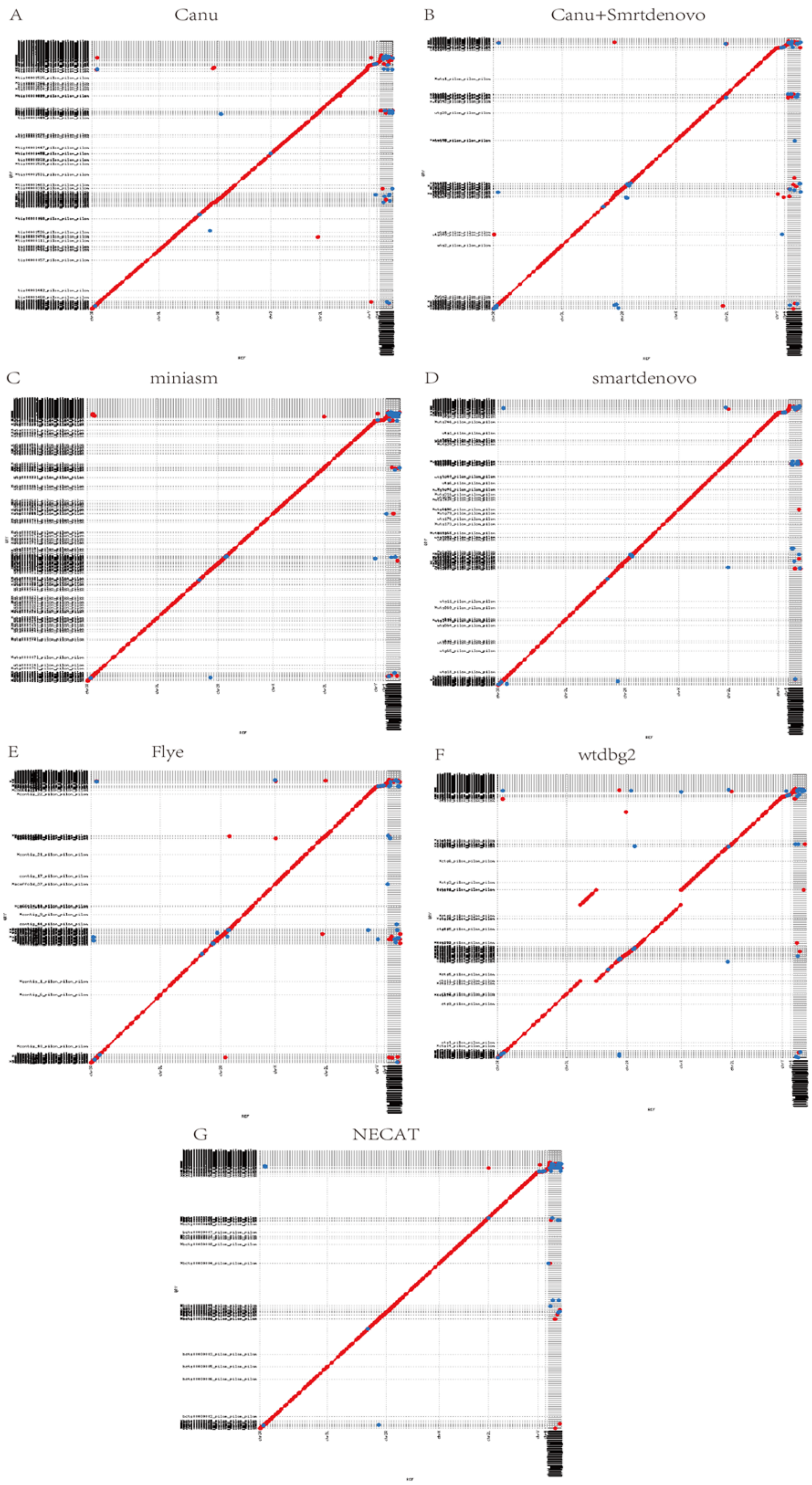


Supplementary Figure 9

Mummerplot of the assembled contig and *D. melanogaster* reference genome.

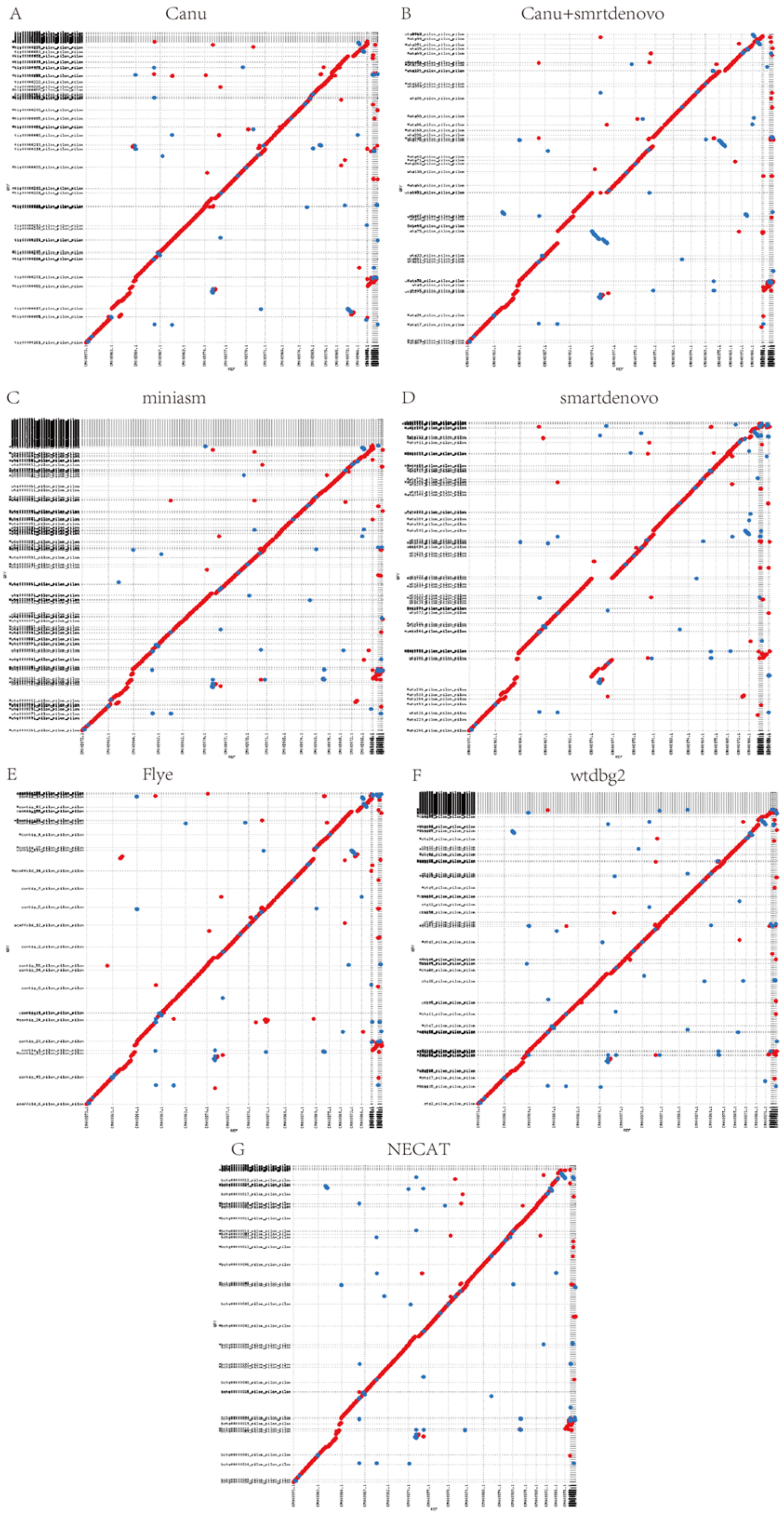


Supplementary Figure 10

Mummerplot of the assembled contig and *C. reinhardtii* reference genome.

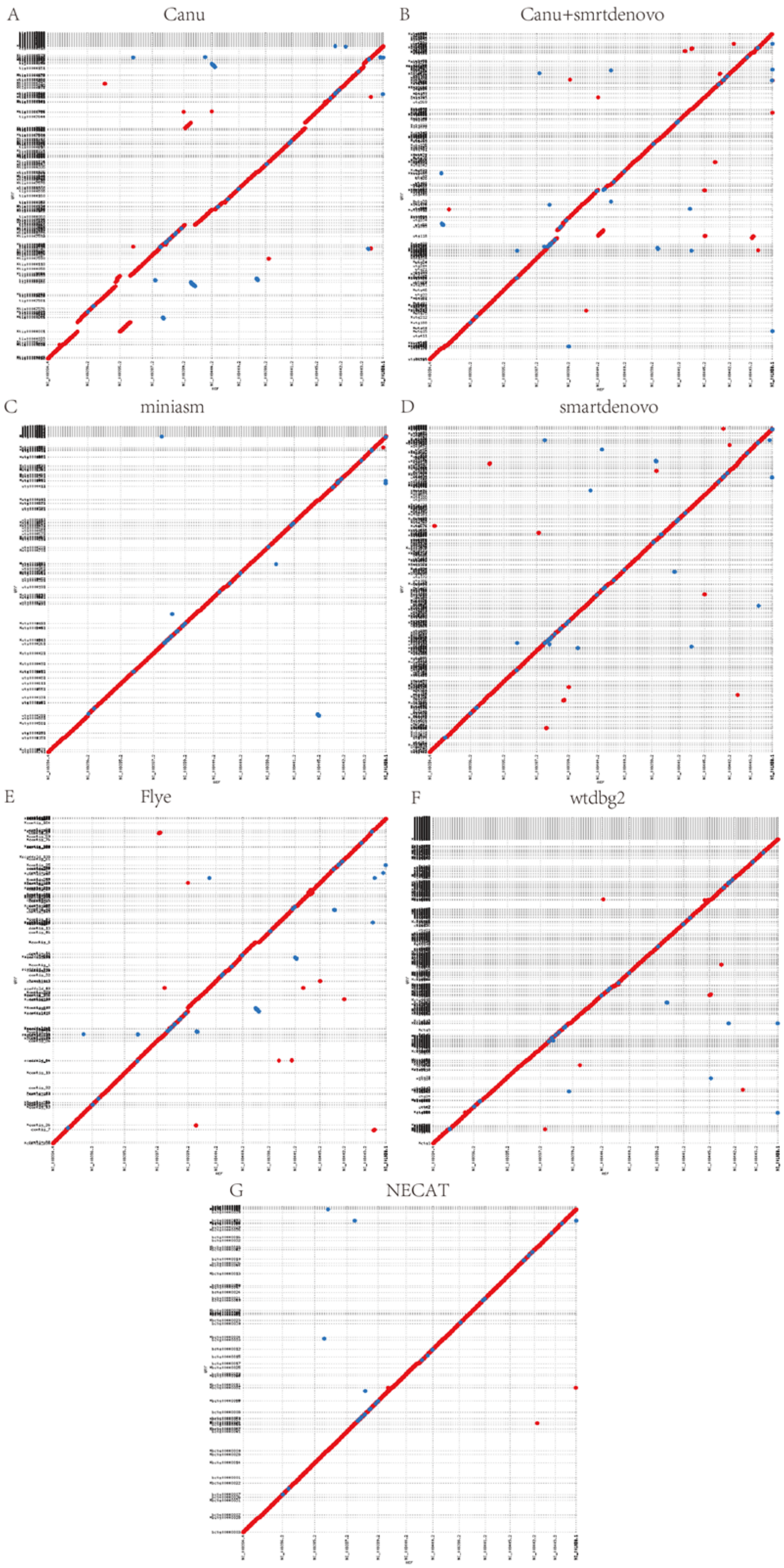


Supplementary Figure 11

Mummerplot of the assembled contig and *O. sativa* reference genome.

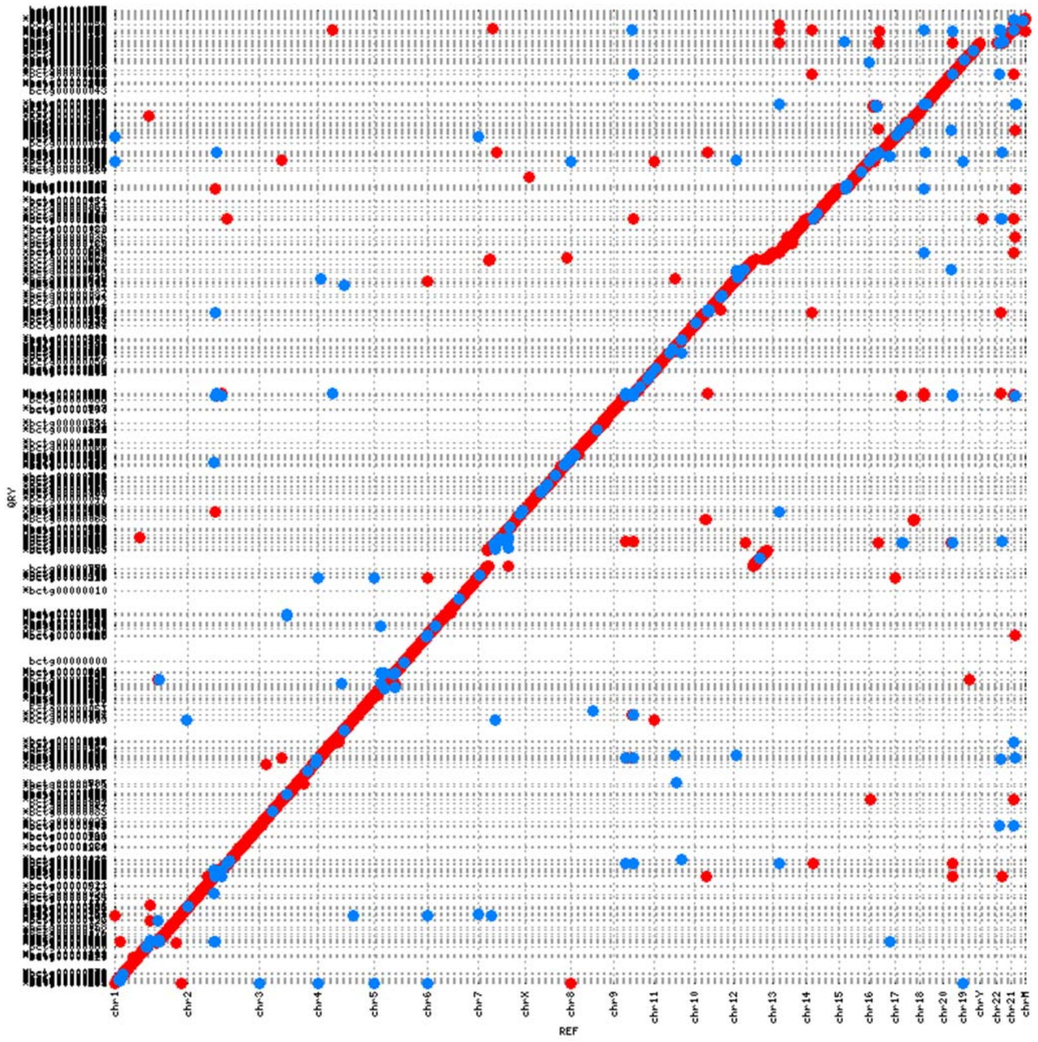


Supplementary Figure 12

Mummerplot of new assembled contigs of NA12878 Nanopore and hg38 reference genome. An alignment dotplot shows the relationship between the contig assembled using Nanopore (y-axis) and GRCh38 reference genome (x-axis). Contig and chromosome boundaries are displayed as dotted lines (horizontal and vertical, respectively).

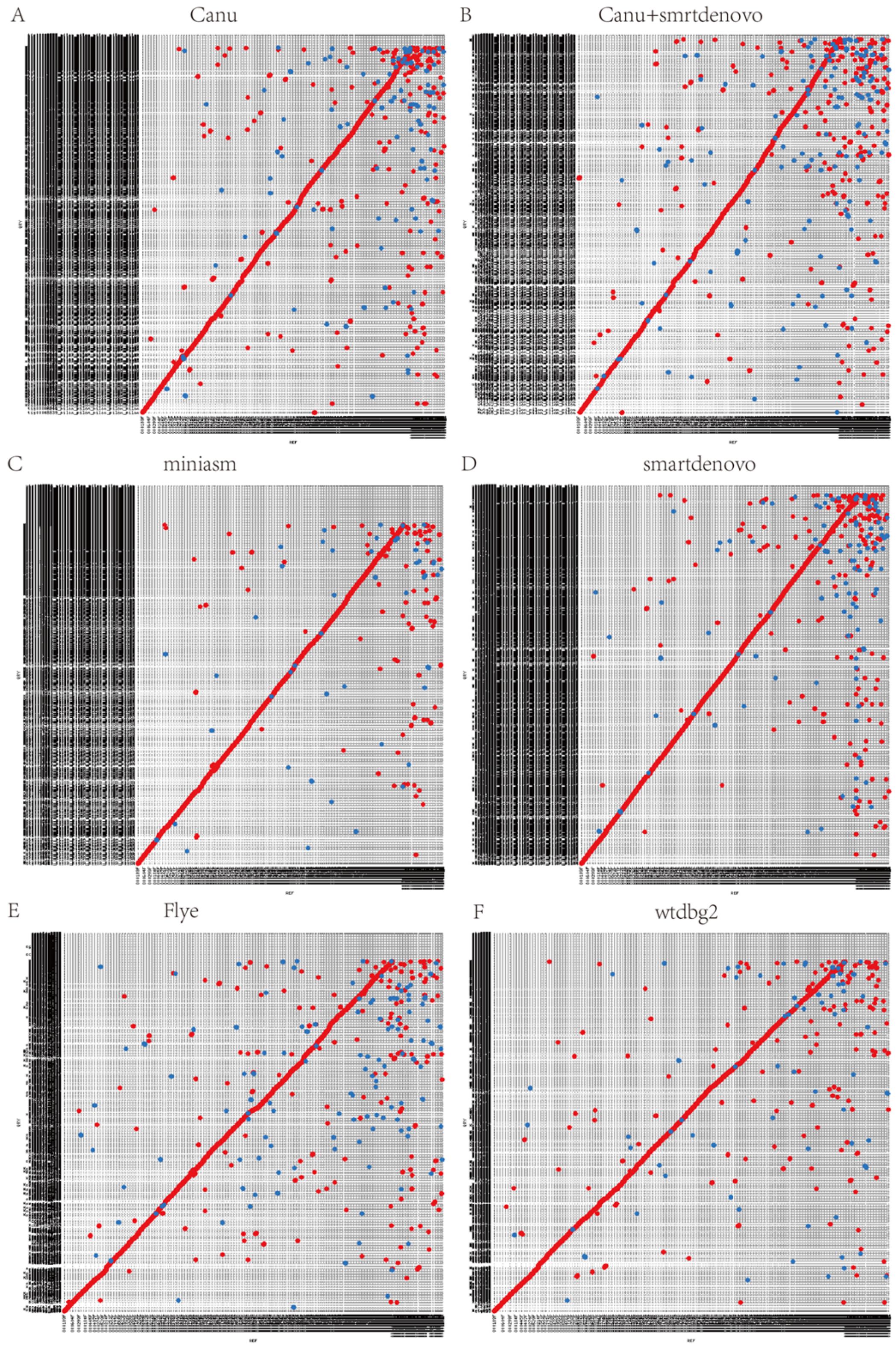


Supplementary Figure 13

Mummerplot of the NECAT contig and *the other assembled* contig from S. pennellii.

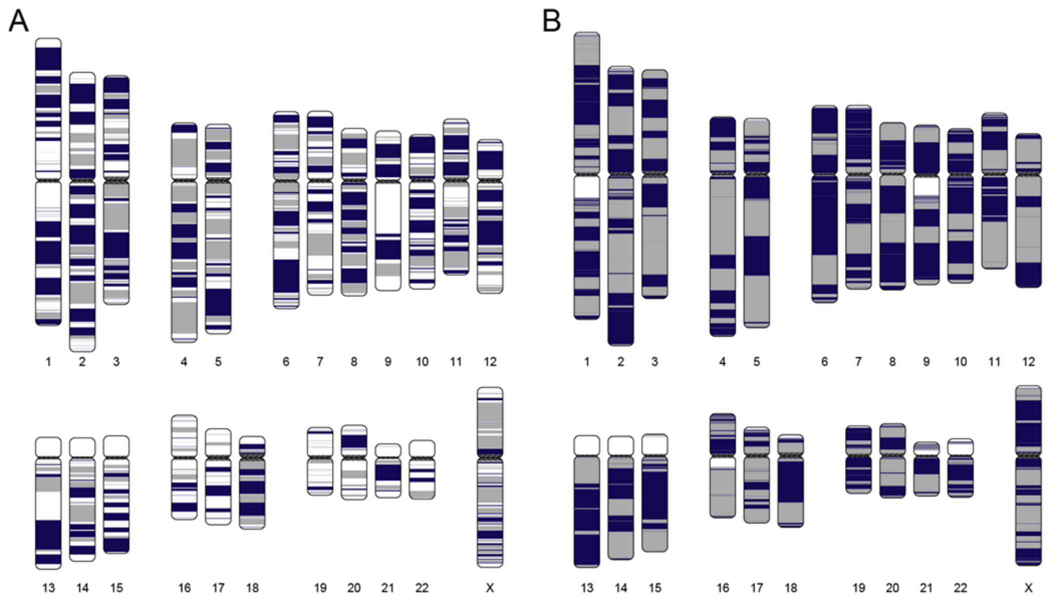


Supplementary Figure 14

Continuity analysis of NECAT and Canu Nanopore *H. sapiens* NA12878 assembly. (A) NECAT assembly. (B) Canu assembly. Human chromosomes are painted with assembled contigs using ColoredChromosomes package. Alternating shades indicate adjacent contigs (each vertical transition from gray to black represents a contig boundary or alignment breakpoint).

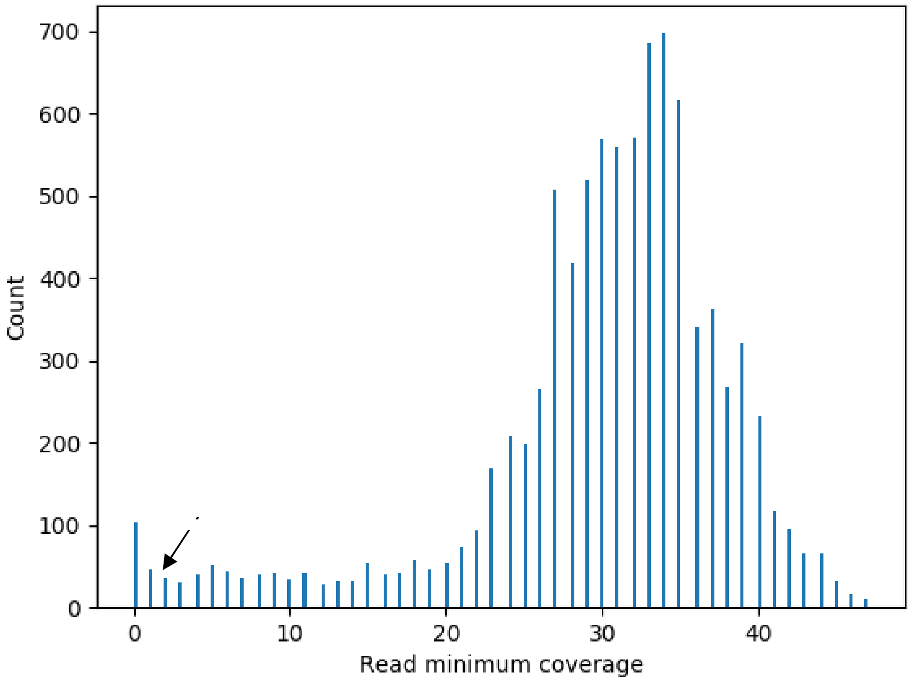


Supplementary Figure 15

Histogram of minimum coverage for the *S. cerevisiae* dataset.
